# De novo E-cadherin/catenin complex formation controls basal epithelial mechanics and force transmission for apoptotic cell clearance

**DOI:** 10.64898/2025.12.08.692667

**Authors:** Hanna-Maria Häkkinen, Marta Batet, Laura F. Bianchi, Senda Jiménez-Delgado, Fabio Pezzano, Stefan Wieser, Luca Ciampa, Esteban Hoijman, Carina Vibe, Sofie Wijma, Verena Ruprecht

**Author notes:** These authors contributed equally (Hanna-Maria Häkkinen, Marta Batet, Laura F. Bianchi).

## Abstract

Beyond serving as cohesive barriers, epithelia clear apoptotic cells to regulate development, homeostasis and inflammation. How epithelial cells remodel their shape during phagocytosis while preserving tissue integrity, and the role of adhesion receptors in this process, remain unclear. Using live in vivo imaging of phagocyte–target interactions in zebrafish (*Danio rerio*) embryos, we show that basal and apical epithelial domains are mechanically decoupled, enabling engulfment without disrupting tissue cohesion. We identify a dynamic assembly of E-cadherin/catenin complexes at the basal epithelial surface in contact with apoptotic cells. Targeted perturbations reveal two critical functions of de novo E-cadherin/catenin complex formation at the phagocytic synapse: α-catenin acts as a physical linker transmitting actin-generated forces required for engulfment, while p120-catenin restrains Myosin II activity, enabling efficient clearance. We further demonstrate the conservation of E-cadherin-dependent apoptotic cell clearance in the mouse trophectoderm. These findings reveal that the E-cadherin/catenin complex is repurposed at the phagocytic synapse as a mechano-regulator of epithelial efferocytosis beyond its canonical role in tissue cohesion.

## Introduction

Epithelia are connected tissues that build structural and functional units relevant to organismal physiology. The ability of epithelial tissues to engulf apoptotic cells identifies them as non-professional phagocytes, a function that is widely conserved across species and plays vital roles in development, tissue homeostasis, repair and the regulation of inflammation^1,2,3,4^. Efferocytosis, the clearance of apoptotic cells, is performed in different epithelial tissues of the adult organism^5,6,7^, and establishes an innate immune function in the developing embryo^8^. The removal of apoptotic cells in the earliest stages of life is of particular relevance considering that both stochastic and programmed events of cell death frequently occur during embryo development^9,10^. An outer squamous epithelium is formed as the first differentiated tissue at blastula stage in species developing both *in utero* and *ex utero*^11,12^, termed enveloping epithelial layer (EVL) in zebrafish^13^ and trophectoderm in mammalian species^14^. We previously showed that the embryonic surface epithelium functions as an efficient innate scavenger tissue for the removal of apoptotic cells in zebrafish and mouse embryos^8^ that is active already prior to embryonic lineage specification.

To guarantee epithelial integrity and function, tissue cohesion needs to be maintained during the phagocytic activity of epithelial cells. Adherens junctions establish stable cell-cell connections between epithelial cells, primarily through the homotypic trans-binding of E-cadherin adhesion receptors^15^. These junctions are mechanically reinforced by interactions between E-cadherin and the underlying actomyosin cortex^16,17^, creating a dynamic interface that couples adhesion sites to cytoskeletal force generation, involving a close mechano-signalling crosstalk^18^.

Phagocytic clearance involves a profound reorganization of the actin cytoskeleton and cellular shape changes during the engulfment of apoptotic targets^19^. This raises the question how dynamic events of cytoskeletal remodelling during phagocytic clearance at the single cell level are coordinated with the mechanical and structural stability of epithelia at the tissue level. Here we combined quantitative live imaging with targeted pharmacological and genetic tools to address the role of cell-cell adhesion in apoptotic cell clearance by epithelial cells in vivo. We identified that E-cadherin is essential for the phagocytic activity of epithelial cells: reducing E-cadherin levels or disrupting its binding capacity leads to a failure of apoptotic cell clearance, prior to affecting epithelial tissue integrity. High-resolution in vivo imaging revealed that E-cadherin at the basal epithelial surface dynamically assembled into a functional signalling complex upon interaction with apoptotic targets, with a local recruitment of α-, β-, and p120-catenin to the phagocytic synapse. We show that de novo formation of the E-cadherin complex regulates apoptotic cell engulfment by two essential mechano-regulatory mechanisms: First, α-catenin mediates mechanical force transmission to ensure proper phagocytic cup progression and apoptotic target engulfment, and abolishing the actin binding domain of α-catenin disrupts apoptotic cell clearance. Second, p120-catenin is required to control Myosin II localization and contractile activity at the phagocytic synapse, and interfering with p120-catenin recruitment inhibits apoptotic target clearance, an effect that can be restored by reducing Myosin II activity via pharmacological or genetic means.

Together, our findings reveal an important physiological function of non-junctional E-cadherin for the phagocytic clearance of apoptotic cells by epithelia. De novo assembly of E-cadherin/catenin complexes at the basal epithelial surface in contact with apoptotic cells regulates basal cortex dynamics and transmits mechanical forces from the actin cytoskeleton for successful apoptotic cell engulfment. Epithelia thereby establish a fundamental mechanism to actively survey their environment and clear dying cells through repurposing the E-cadherin/catenin signalling complex at the phagocytic synapse outside cell-cell contacts, while maintaining tissue cohesion and barrier integrity.

## Results

### Epithelial tissue architecture is maintained during apoptotic cell clearance

To study the spatio-temporal clearance dynamics of apoptotic cells by epithelia in vivo, we employed a previously established ‘apoptotic stress induction’ assay in the zebrafish embryo^8^. Cell death was triggered via mosaic co-expression of the pro-apoptotic protein Bax in combination with a fluorescently tagged membrane marker (Lyn-tdTomato; Bax+ cells), leading to apoptotic cell death in a sub-population of embryonic progenitor cells during blastula stage (Figure 1A). Dynamic surface protrusions facilitate a continuous probing of the tissue environment and are involved in the phagocytic clearance activity of professional immune cells^20^. Membrane ruffles have also been documented at the basal surface of epithelial cells^21,22^. Live cell in vivo imaging revealed that epithelial cells showed highly dynamic membrane protrusions on their basal surface (Figure 1B, Supplementary Movie 1), independently of the presence of apoptotic cell targets. In comparison, we observed slower junctional dynamics at the apical site (Figure 1B, Supplementary Movie 1), as quantified in dynamic flow maps of the basal versus apical epithelial regions (Figure 1B). In vivo imaging of basal epithelial cell dynamics further enabled us to visualize the interaction of ruffles with apoptotic cells and to capture individual uptake events followed by the formation of phagosomes (Figure 1C; Supplementary Movie 2). Notably, the apical epithelial cell area was maintained during the phagocytic uptake of apoptotic cells (Figure 1C,D), and epithelial cells in contact with apoptotic cells or live cells had a similar apical cell area (Supplementary Figure 1A). These data support that the morphodynamic remodelling of the basal epithelial surface during phagocytosis does not influence apical cell and tissue architecture, suggesting that basal and apical epithelial cell surface dynamics are mechanically decoupled. Together, these findings reveal that the basal epithelial surface is a highly dynamic interface, enabling epithelia, like professional immune cells, to extend their surface and actively survey the local tissue environment, while maintaining tissue cohesion.

**Figure 1.**
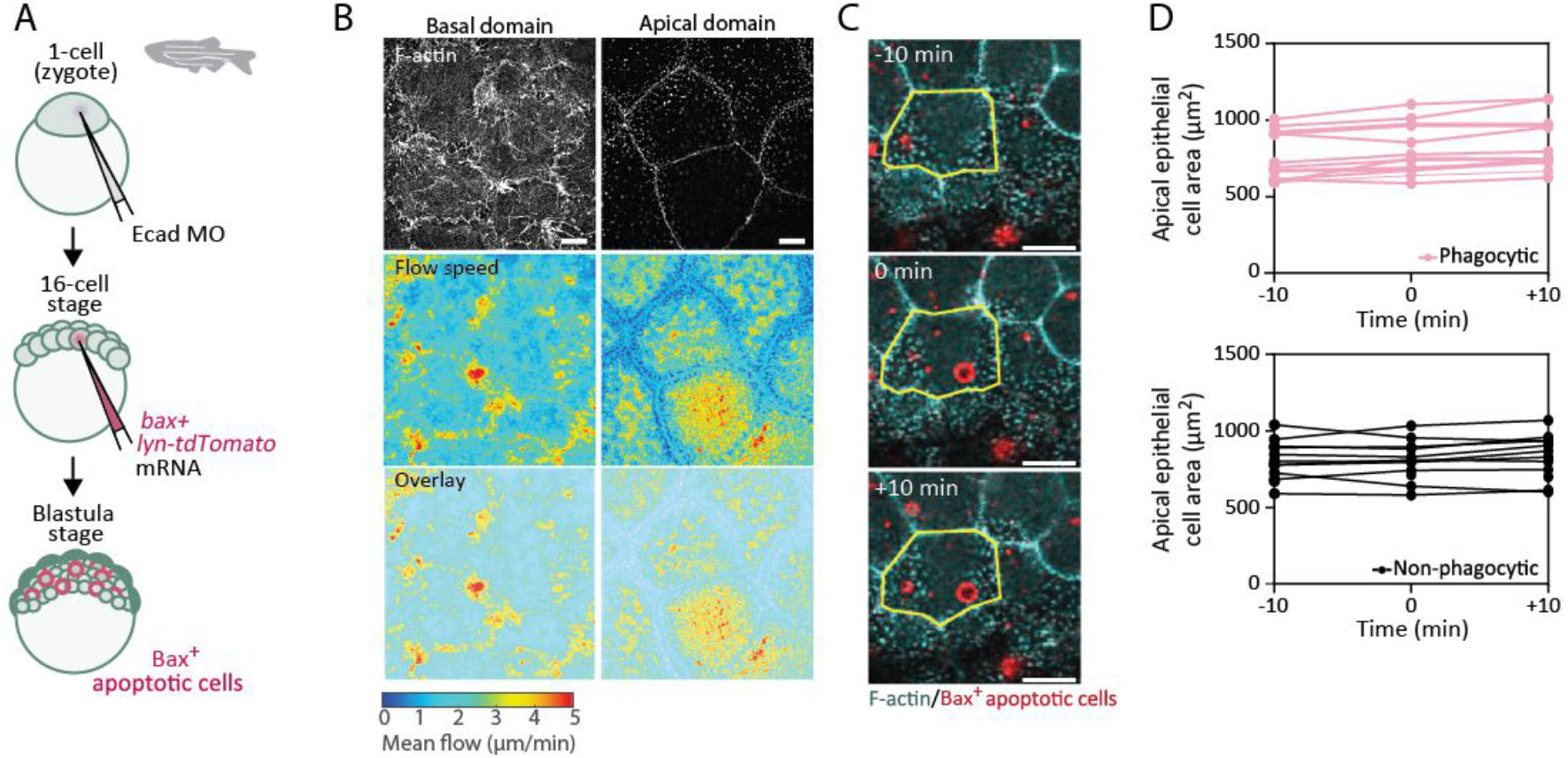
Epithelial tissue architecture is maintained during apoptotic cell clearance. **A)** Sketch of the experimental setup showing embryo microinjection at 1-cell (zygote) stage with E-cadherin morpholino (Ecad MO) for the ubiquitous down-regulation of E-cadherin (top) and single blastomere mRNA injection at 16-cell stage for mosaic expression (middle) to induce apoptosis in a subpopulation of cells (Bax+ cells, bottom) via the co-expression of a pro-apoptotic factor (Bax) and a plasma membrane (PM) marker (red, Lyn-tdTomato) in the embryo blastula. **B)** Representative images of F-actin localization at the basal and apical epithelial domains (top), flow speed maps of F-actin dynamics (middle) and their overlay (bottom). **C-D)** Representative dual-colour fluorescence images (overlay) of the apical epithelial area (yellow lines) 10 min before, during and 10 min after an uptake of an apoptotic target (Bax+ cells, red) and its quantification in D) (top, n=18 cells, 4 embryos) in comparison with neighbouring non-phagocytic epithelial cells (bottom; n=12 cells, 4 embryos). All embryos were obtained from the transgenic Tg(actb2:Lifeact-GFP) line. Scale bars: 10 μm (B) and 20 μm (C).

### The basal epithelial E-cadherin pool is required for apoptotic cell clearance

Adherens junctions are integral to epithelial tissue cohesion and apico-basal cell polarity^23^. 3D SIM imaging using the KI(mlanYFP)^xt17^cdh1-YFP line (E-cadherin-YFP), in which a fluorescent tag was introduced at the endogenous locus of E-cadherin^24^, confirmed the baso-lateral localization of E-cadherin (Supplementary Figure 1B). Notably, we further detected the presence of E-cadherin at the phagocytic synapse (Figure 2A; Supplementary Movie 3), and its transient enrichment on newly formed phagosomes (Supplementary Figure 1C; Supplementary Movie 4). We previously found that both phagocytic cups and ‘epithelial arms’ are involved in mediating a cooperative apoptotic cell clearance by epithelial cells^8^. Notably, epithelial arms are a specific type of basal epithelial protrusions, which we identified can generate mechanical pushing forces on apoptotic targets and drive their movement and dispersal at the tissue level, without the requirement for cell autonomous motility in apoptotic cells^8^. Live cell in vivo imaging of E-cadherin-YFP revealed the localization of E-cadherin both at phagocytic cups and at ‘epithelial arm’ protrusions associated to an apoptotic target (Figure 2A, Supplementary Figure 1 C,D; Supplementary Movie 5). Moreover, the junctional localization of E-cadherin was maintained in epithelial cells with apoptotic cargo (Supplementary Figure 1E). These observations suggest an involvement of non-junctional E-cadherin at the basal epithelial domain in the phagocytic clearance of apoptotic cells.

**Figure 2.**
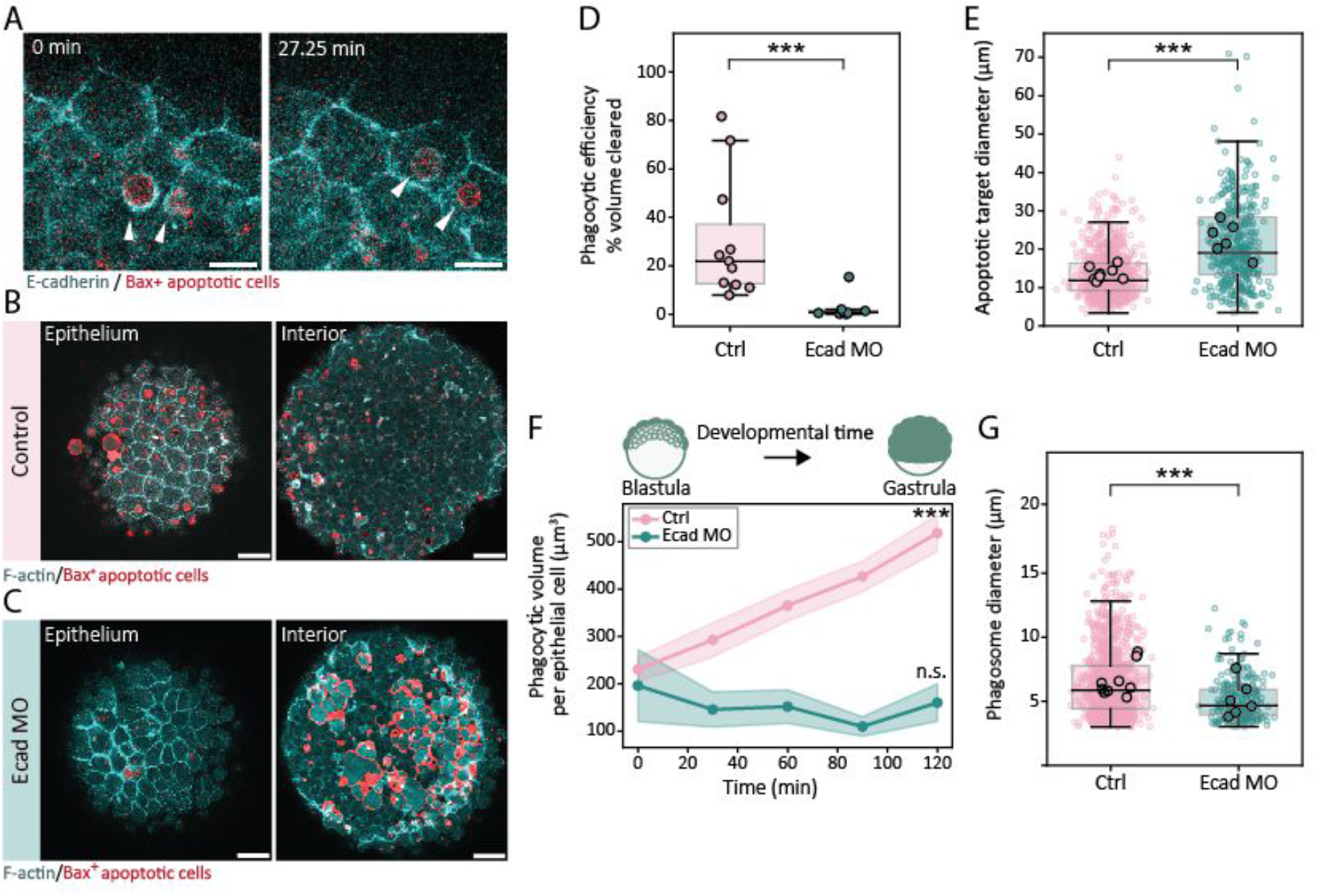
E-cadherin is necessary for epithelial efferocytosis. **A)** Representative dual-colour fluorescence images (overlay) of E-cadherin localization (cyan) at two different phagocytic cups (arrowheads). **B-D)** Representative dual-colour fluorescence images (B, C) and their quantification (D) showing the location of apoptotic targets (Bax+ cells, red) in the epithelium (left) and the embryo interior (right) at shield stage (6 hpf) in control (B) and Ecad MO embryos (C). Phagocytic clearance efficiency was measured in control (n=11 embryos) and Ecad MO (n=8 embryos) embryos at shield stage (6.5 hpf). N=4. Mann-Whitney test, p=3*10^-4^. **E)** Diameter of apoptotic targets in the embryo interior at shield stage (6.5 hpf; n=855 targets, 9 embryos (Ctrl); n=394 targets, 6 embryos (Ecad MO); N=3. Manb-Whitney test, p<10^-4^. **F)** Total volume of phagosomes per epithelial cell from blastula to gastrula stage (4.5-6.5 hpf; Ctrl: n=76 (0 min), n=147 (30 min), n=173 (60 min), n=202 (90 min), n=226 (120 min) epithelial cells; Ecad MO: n=8 (0 min), n=15 (30 min), n=17 (60 min), n=25 (90 min), n=31 (120 min) epithelial cells; 3 embryos per condition; N=3) Data points and shaded areas represent the mean+/-SEM. Kruskal-Wallis test (start-to-endpoint), Ctrl: p<10^-4^; Ecad MO: p=0.6048. **G)** Diameter of phagosomes in the epithelium at shield stage (6.5 hpf; n=1179 phagosomes from 9 embryos (Ctrl), n=253 phagosomes from 6 embryos (Ecad MO); N=3). Mann-Whitney test, p=10^-4^. Embryos were obtained from the transgenic KI(mlanYFP)^xt17^cdh1-YFP line (A) and Tg(actb2:Lifeact-GFP) line (B,C). Large dots each represent an embryo (D) or the mean value for each embryo (E,F). N indicates the number of independent experiments. Box plots show the median, the 1st and 3rd quartile and the 1.5x inter-quartile range. All statistical tests are two-sided. Scale bars: 10 μm (A) and 50 μm (B,C).

To directly assess the role of E-cadherin in the phagocytic clearance process of apoptotic cells in vivo, we targeted *cdh1* expression, which encodes E-cadherin, representing the major cadherin isoform expressed in the embryonic epithelium^8,25^. Induction of apoptosis was followed by the rapid clearance of dying cells by the outer surface epithelium of the embryo within 7 hours post fertilization (hpf; Figure 2B; Supplementary Movie 6), as described previously^8^. In contrast, morpholino (MO) interference with *cdh1* expression, using a validated MO sequence^27^, caused a pronounced defect in phagocytic clearance capacity compared to control embryos (Figure 2C,D; Supplementary Movie 6). Quantitative immunofluorescence and live imaging of E-cadherin-YFP, together with Western Blot analysis, confirmed a reduction of E-cadherin levels in the embryonic epithelium (Supplementary Figure 2A-E). The clearance defect in E-cadherin deficient epithelial cells was accompanied by an increased size of apoptotic targets and their accumulation in the embryo interior (Figure 2E). In addition, the rate of phagosome formation was substantially reduced, measured by the total phagosome volume per epithelial cell over time from blastula to gastrula stage (4.5-6.5 hpf) (Figure 2F; Supplementary Movie 7). Moreover, the phagosome size in epithelial cells with a residual uptake in E-cadherin morphant embryos was significantly smaller compared to control embryos (Figure 2G, Supplementary Figure 2F), consistent with an overall defect in tissue clearance capacity upon E-cadherin depletion.

At the single epithelial cell level, F-actin remodelling was initiated upon contact with apoptotic cells both in control and E-cadherin morphant conditions (Supplementary Figure 2G,H and Supplementary Movie 8). However, E-cadherin reduction compromised the formation of functional phagocytic cups and the frequency of epithelial ‘arms’ was strongly decreased (Supplementary Figure 2I,J). In addition, the average and maximum apoptotic cell speed in vivo due to residual epithelial ‘arm’ protrusions was reduced, associated with a restricted spatial dispersal of apoptotic targets compared to control conditions (Supplementary Figure 2K-N, Supplementary Movie 9). These results indicate a direct requirement of E-cadherin for apoptotic cell clearance by regulating basal epithelial protrusion dynamics that mediate target interaction and uptake.

To exclude that the clearance defect in E-cadherin deficient embryos was caused by an alteration in epithelial tissue specification, we validated the expression of epithelial-specific genes (*chuk*, *irf6*, *krt18a1* and *krt4*) by qPCR analysis. Comparison of expression levels at two developmental time points (sphere and shield stage; 4/6 hpf) revealed no significant changes in E-cadherin deficient versus control embryos (Supplementary Figure 3A). Tissue polarity upon E-cadherin reduction was further intact as shown by the localization of aPKC on the apical area of the epithelium at blastula and later gastrulation stages (4/6/8 hpf; Supplementary Figure 3B). In addition, characteristic actin distributions were maintained in E-cadherin deficient embryos, such as apical microridges and junctional actin pools (Supplementary Figure 3C), and E-cadherin retained its localization at the basolateral domain, albeit at reduced levels (Supplementary Figure 3D). Moreover, while E-cadherin deficient embryos showed a typical delay of gastrulation movements as described previously^27^, morphological characteristics of the epithelium were maintained in control and E-cadherin deficient embryos, including epithelial cell shape and size (4/6/8 hpf; Supplementary Figure 3E,F).

We next probed the involvement of junctional versus basal E-cadherin in the clearance process. For this purpose, we acutely inhibited the basal E-cadherin pool using tissue microinjections of a specific blocking antibody (DECMA-1) targeted against the extracellular EC1 domain of E-cadherin. We validated that epithelial cell polarity and adherens junctions were maintained upon DECMA1 injection (Supplementary Figure 3G). Microinjections of DECMA-1 and DMEM (control) into the embryo blastula cap were conducted at two different developmental time points. First, DECMA-1 was injected prior to the start of phagocytic tissue clearance (early dome stage (4.5 hpf), Figure 3A). DECMA-1 injection strongly reduced tissue clearance, as monitored over a duration of 120 min after injection (Figure 3B,C; Supplementary Movie 10), corroborating our observations in E-cadherin morphant embryos. Phagosome sizes were further decreased (Figure 3D) and the apoptotic target diameter in the embryo interior was enlarged (Figure 3E), similar to observations in E-cadherin morphant embryos. We next performed injections of DECMA-1 at a time point when phagocytic clearance had already commenced and apoptotic cell size was reduced (50% epiboly stage (5.5 hpf), Supplementary Figure 3H,I). While the phagocytic uptake of apoptotic cells by the epithelium progressively increased in the control condition, phagocytic clearance was stalled in embryos upon DECMA-1 injection over the measurement period of 120 min (Figure 3F; Supplementary Movie 11). Together, these data support that the E-cadherin pool at the basal epithelial surface is required for the clearance of both large and fragmented apoptotic targets.

**Figure 3.**
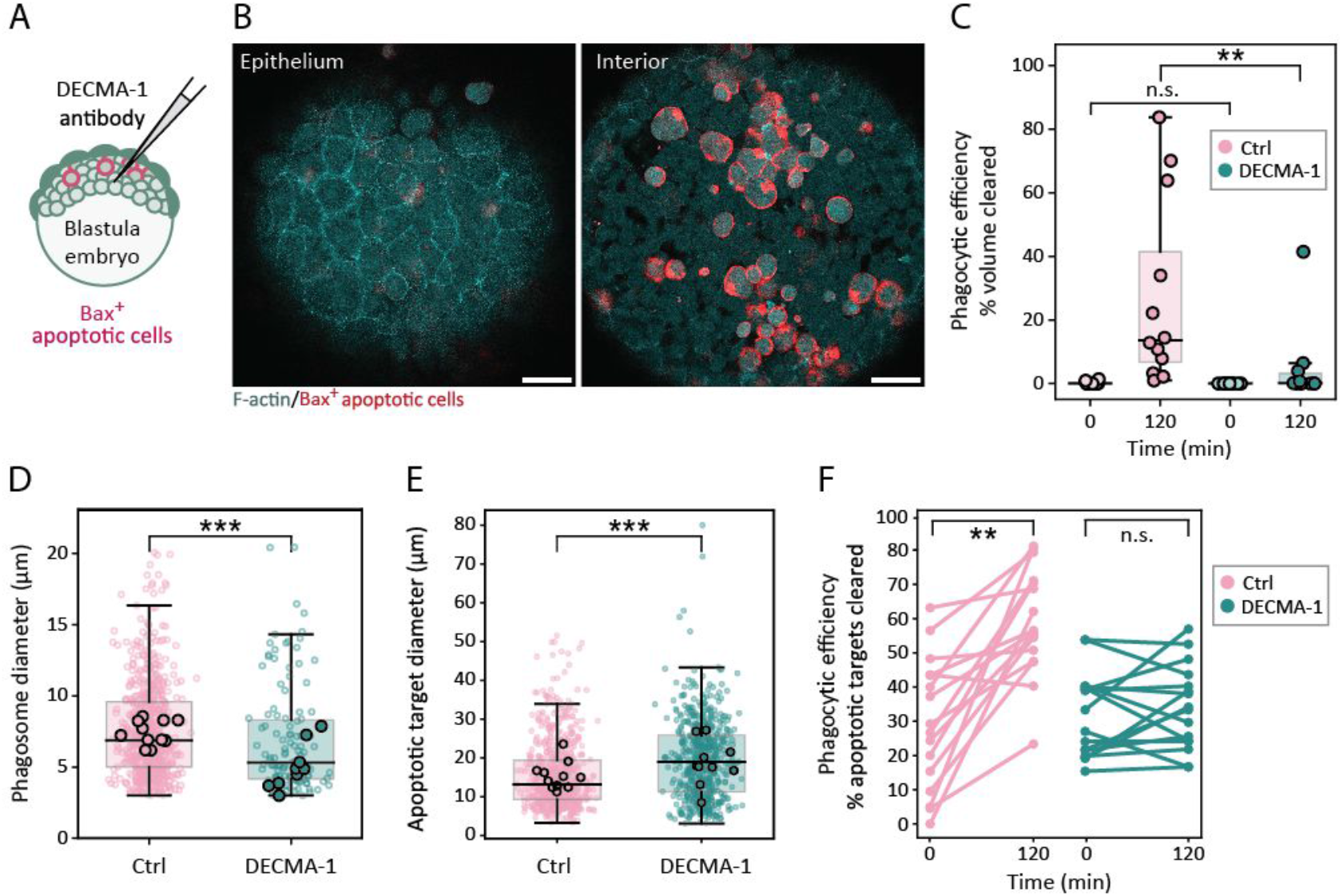
Acute perturbation of E-cadherin ectodomain binding leads to a rapid stalling of apoptotic tissue clearance. **A)** Experimental sketch representing the acute inhibition of E-cadherin binding by microinjection of DECMA-1 blocking antibody into the embryo blastula cap. **B)** Representative dual-colour fluorescence images (overlay) showing the location of apoptotic targets (Bax+ cells, red) in the epithelium (left) and the embryo interior (right) at shield stage (6 hpf) in DECMA-1 injected embryos expressing Lifeact-GFP (cyan). **C)** Phagocytic clearance efficiency (volume fraction) in DECMA-1 (n=10 embryos) and DMEM medium (Ctrl, n=12 embryos) injected embryos at dome stage (4.5 hpf) and analysed for phagocytic uptake at dome stage (4.5 hpf, t=0 min) and at shield stage (6.5 hpf, t=120 min). N=3. Mann-Whitney test: p=0.2209 (0 min), p=0.0020 (120 min). **D)** Diameter of individual phagosomes in control (Ctrl, n=631 phagosomes from 12 embryos) and DECMA-1 injected embryos (n=123 phagosomes from 10 embryos) at shield stage (6.5 hpf). N=3. Mann-Whitney test, p=0.0002. **E)** Diameter of individual apoptotic cell targets in DECMA-1 (n=571 targets from 10 embryos) and DMEM medium (Ctrl, n=715 targets from 12 embryos) injected embryos at shield stage (6.5 hpf). N=3. Mann-Whiteny test, p<10^-4^. **F)** Phagocytic clearance efficiency derived as the percentage of targets engulfed by the epithelium from the total number of available apoptotic targets (with maximum target diameter ≤ 8 μm) in control embryos (n=16) and embryos injected with DECMA-1 blocking antibody (n=16) at 50% epiboly stage (5.5 hpf) and analysed for phagocytic uptake at 50% epiboly stage (5.5 hpf, t=0 min) and 80% epiboly (8 hpf, t=120 min). N=3. Data points for individual embryos are connected over time. Paired t-test, p<10^-4^ (Ctrl), p=0.4173 (DECMA-1). Embryos were obtained from the transgenic Tg(actb2:Lifeact-GFP) line. Large dots represent an embryo (C) or the mean value for each embryo (D,E). N indicates the number of independent experiments. Box plots show the median, the 1st and 3rd quartile and the 1.5x inter-quartile range. All statistical tests are two-sided. Scale bars: 50 μm (B).

### E-cadherin trans-binding is dispensable at the phagocytic synapse

To assess whether E-cadherin trans-binding is required at the phagocytic synapse for target interaction and uptake, we next transplanted apoptotic cells obtained from control or E-cadherin deficient donor embryos (Ecad MO) into wild type host embryos (Figure 4A). Quantification of the phagocytic clearance efficiency showed no significant change between these conditions (Figure 4B,C; Supplementary Movie 12), similarly to the mean phagosome size (Figure 4D). Furthermore, no difference in the spatial spreading of apoptotic targets due to the activity of epithelial arm protrusions was detected (Supplementary Figure 4A,B), with a similar mean and maximum apoptotic target speed (Supplementary Figure 4C,D).

**Figure 4.**
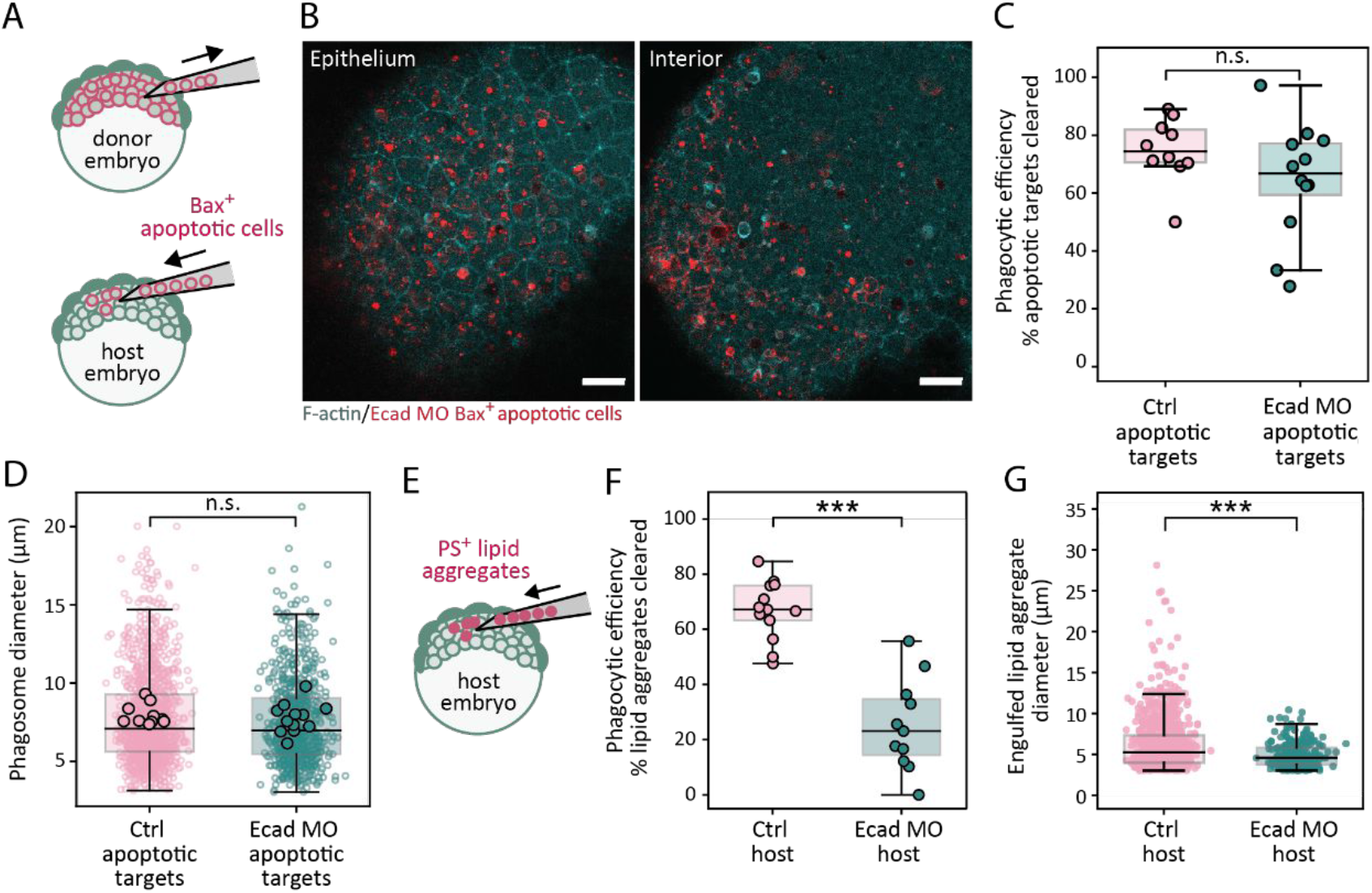
Homotypic E-cadherin trans-binding is dispensable for phagocyte-target interactions and engulfment. **A)** Experimental sketch of the transplantation assay to transfer apoptotic targets from a donor embryo (control or E-cadherin morphant) into a host embryo at blastula stage (4 hpf) prior to imaging. **B)** Representative dual-colour fluorescence images of the surface epithelium (left) and embryo interior (right) after transplantation of Ecad MO targets (red) in an embryo expressing Lifeact-GFP (cyan) at shield stage (6.5 hpf). **C)** Phagocytic clearance efficiency derived as the percentage of apoptotic targets cleared by the embryonic epithelium in embryos transplanted with apoptotic cells from control donor embryos (Ctrl, n=10) and apoptotic cells from Ecad MO donor embryos (Ecad MO, n=12). N=5. Unpaired t-test, p=0.1573. **D)** Diameter of phagosomes cleared by the epithelium in embryos transplanted with control apoptotic cells (Ctrl, n=1130 phagosomes from 10 embryos) and apoptotic cells obtained from Ecad MO embryos (Ecad MO, n=702 phagosomes from 12 embryos). N=5. Mann-Whitney test, p=0.1991. **E)** Experimental sketch of the transplantation assay to transfer PS^+^ lipid aggregates into the blastula of control or E-cadherin MO host embryos at sphere stage (4 hpf). **F)** Phagocytic clearance efficiency derived as the percentage of PS^+^ lipid aggregates engulfed by the epithelium from the total amount of aggregates present in Ctrl (n=13) and Ecad MO (n=10) conditions. N=3. Mann-Whitney test, p<10^-4^. **G)** Phagosome diameter of cleared PS^+^ lipid aggregates in control (n=892 phagosomes from 13 embryos) and Ecad MO embryos (n=277 phagosomes from 10 embryos) at 75% epiboly stage (8 hpf). N=3. Each dot represents a phagosome. Mann-Whitney test, p<10^-4^. All embryos were obtained from the Tg(actb2:Lifeact-GFP) line. Large dots represent an embryo (C,F) or the mean value for each embryo (D). N indicates the number of independent experiments. Box plots show the median, the 1st and 3rd quartile and the 1.5x inter-quartile range. All statistical tests are two-sided. Scale bars: 50 μm (B).

To further validate that E-cadherin trans-binding was dispensable at the phagocytic synapse between epithelial and apoptotic cells, we next transplanted synthetic targets that mimic apoptotic cells. For this purpose, we generated lipid aggregates containing the apoptotic cell recognition factor phosphatidylserine (PS) (Figure 4E). Transplantation of synthetic targets into wild type embryos confirmed their rapid uptake by the epithelium via the formation of F-actin enriched phagocytic cups similar to apoptotic cells (Figure 4F; Supplementary Figure 4E,F; Supplementary Movie 13). Upon engulfment of lipid aggregates, phagosomes with mean sizes comparable to the uptake of apoptotic cells were formed (Figure 4G and Figure 2G). In addition, E-cadherin enriched around lipid aggregates during the interaction and engulfment of synthetic targets by epithelial cells (Supplementary Figure 4G; Supplementary Movie 14). Notably, in E-cadherin deficient embryos synthetic targets were not uptaken and remained in the embryo interior (Figure 4F; Supplementary Figure 4F; Supplementary Movie 13), consistent with the clearance defect of apoptotic cells in epithelial cells with reduced E-cadherin levels. Together, these data support that E-cadherin is essential for apoptotic cell clearance by a mechanism independent of homotypic E-cadherin trans-binding at the phagocyte-target interface.

### De novo assembly of the E-cadherin/catenin complex at the phagocytic synapse on the basal epithelial cell surface

To dissect how E-cadherin regulates apoptotic cell clearance, we next studied its interaction with canonical binding partners. α- and β-catenin are core components of the E-cadherin complex, linking adhesion receptors to the actin cytoskeleton and coordinating mechanical force transmission^28,29^. To exclusively visualize α- and β-catenin dynamics at the basal epithelial cortex in contact with apoptotic cells, we performed transplantations of apoptotic targets from a donor embryo co-expressing Bax and a membrane marker (Lyn-tdTomato/Cerulean), into a host embryo expressing α-catenin-mCherry or EGFP-β-catenin. In the absence of apoptotic targets, α-catenin localization was not detected at the basal epithelial surface, while it was enriched at cell-cell junctions (Supplementary Figure 5A). Notably, we observed the dynamic recruitment of α-catenin during the formation of phagocytic cups upon apoptotic cell interactions, with a gradual enrichment during phagocytic cup progression (Figure 5A; Supplementary Movie 15). In addition, α-catenin was present at the surface of newly formed phagosomes, from which it was gradually depleted after a few minutes (Figure 5A; Supplementary Movie 15). α-catenin recruitment further occurred in an E-cadherin dependent manner (Supplementary Figure 5B). Similarly, β-catenin was recruited to phagocytic contact sites upon apoptotic target encounter and localized on early phagosomes (Supplementary Figure 5C; Supplementary Movie 16), while it was absent from the basal epithelial surface when no apoptotic target interactions occurred (Supplementary Figure 5D). Moreover, β-catenin retained its junctional localization and did not translocate to the nucleus in control and E-cadherin deficient embryos in epithelial cells that had engulfed phagocytic cargo (Supplementary Figure 5E).

**Figure 5.**
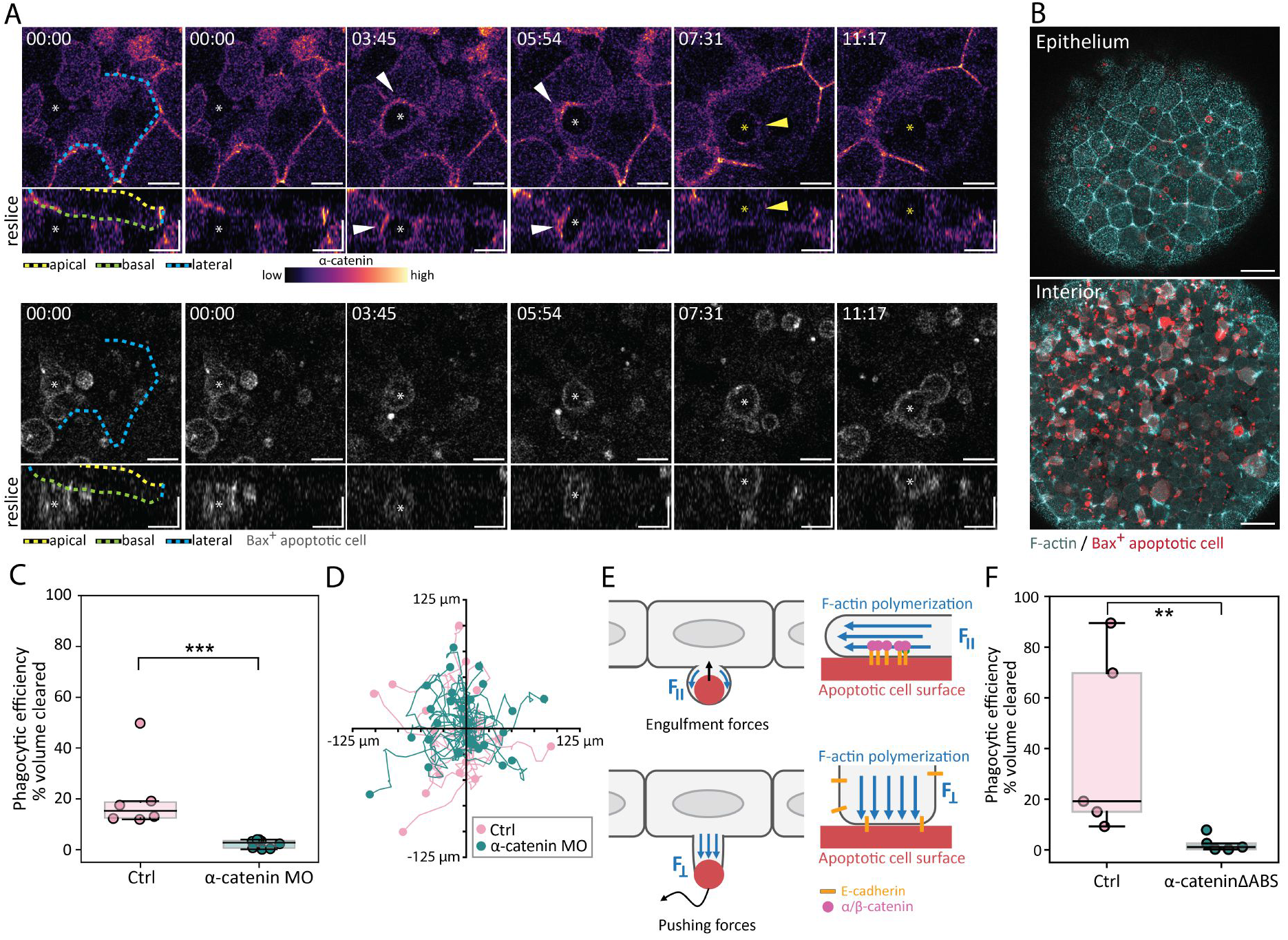
Recruitment of α-catenin to the phagocytic synapse is required for mechanical force transmission during phagocytosis. **A)** Time lapse images showing the dynamic localization of α-catenin-mCherry in epithelial cells (top) during the uptake (white arrowheads) of transplanted apoptotic cells (Bax+ cells, bottom, asterisks) and at a newly formed phagosome (yellow arrowheads). Dashed lines indicate different epithelial surfaces (apical, basal, lateral). Time indicated in min:s. **B)** Dual-colour fluorescence images (overlay) showing the location of apoptotic targets (Bax+ cells, red) in the epithelium (top) and embryo interior (bottom) at shield stage (6.5 hpf) in an α-catenin morphant embryo expressing Lifeact-GFP (cyan). **C)** Quantification of the phagocytic efficiency in control (Ctrl, n=6) and α-catenin deficient embryos (α-catenin MO, n=8) at shield stage (6.5 hpf). N=3. Mann-Whitney test, p=0.0007. **D)** x/y tracks of individual apoptotic targets showing the path travelled over a period of 30 min in control (red, n=17 tracks from 3 embryos) and α catenin MO embryos (cyan, n=27 tracks from 4 embryos). Tracks were centred to the origin. Dots represent endpoint positions. **E)** Model sketch illustrating the role of the E-cadherin/catenin complex in epithelial protrusion formation and force transmission upon interaction with apoptotic cells: During phagocytic uptake (top), the E-cadherin/catenin complex links the surface of the target to the F-actin cytoskeleton via α-catenin, allowing for the transmission of actin polymerization forces tangentially to the target’s surface (F_II_) and phagocytic cup progression. In contrast, the formation of epithelial ‘arm’ protrusions (bottom) is based on F-actin polymerization forces acting normally to the contact site (F_⊥_), which do not depend on α-catenin mediated mechanical coupling to actin for the generation of pushing forces. **F)** Quantification of phagocytic efficiency (volume fraction) for control (Ctrl, n=5) and α-cateninΔABS overexpressing embryos (n=5) at 7.5 hpf. N=2. Mann-Whitney test, p=0.008. Embryos were obtained from WT (A) and the transgenic Tg(actb2:Lifeact-GFP) line (B). Large dots represent an embryo (C,F). N indicates the number of independent experiments. Box plots show the median, the 1st and 3rd quartile and the 1.5x inter-quartile range. All statistical tests are two-sided. Scale bars x: 10 μm, y: 10 μm (A); 40 μm (B).

Together, these observations support the de novo formation of E-cadherin/catenin complexes at the basal epithelial cell surface upon contact with apoptotic targets.

### α-catenin mediates mechanical force transmission for apoptotic cell clearance

Given the important function of α-catenin as a direct physical linker to the actin cytoskeleton, we next assessed its functional role in apoptotic cell clearance. α-catenin connects E-cadherin to the actin cytoskeleton via binding to both β-catenin and F-actin^18^. We thus hypothesized that the functional role of α-catenin in apoptotic cell clearance is associated to the mechanical transmission of actin polymerisation forces needed for phagocytic cup progression. Depleting α-catenin, by using a previously validated morpholino interference approach^30^, led to a pronounced failure in phagocytic cup formation and resulted in a tissue-level defect in phagocytic clearance (Figure 5B,C; Supplementary Movie 17). Downregulation of α-catenin did not affect junctional or basal E-cadherin localization (Supplementary Figure 5F,G), suggesting a direct involvement of α-catenin in the phagocytic uptake process. Of note, the formation of epithelial ‘arms’, which generate mechanical pushing forces on apoptotic targets, remained fully functional in α-catenin deficient epithelial cells and led to apoptotic cell movements and their spatial dispersal similar to control embryos (Figure 5D, Supplementary Figure 5H-J; Supplementary Movie 18).

These results support that the generation of traction forces along the target surface, required for phagocytic cup progression and target engulfment, is compromised in the absence of α-catenin (Figure 5E). By contrast, epithelial arm extensions depend on actin polymerization in the direction normal to the contact interface, independently of tangential traction forces, and thus do not require α-catenin-mediated coupling (Figure 5E).

To directly test the role of α-catenin in mechanical force transmission, we generated a α-catenin mutant construct (α-cateninΔABS) that lacks the actin binding domain^28^. Notably, overexpression of this mutant construct severely compromised apoptotic cell clearance (Figure 5F, Supplementary Movie 19), supporting that direct binding to actin is required for apoptotic cell clearance to take place. Consistently, the formation of pushing protrusions was unaffected (Supplementary Movie 19), supporting that target recognition remains functional, while force transmission required for the progression of phagocytic cups and apoptotic target engulfment failed.

Overall, our results support that the E-cadherin/catenin complex is functionally required at the contact site between epithelial and apoptotic cells during phagocytosis. Specifically, α-catenin establishes the physical coupling necessary for the transmission of actin polymerization forces, essential for the advancement of membrane protrusions during phagocytic cup progression and target engulfment.

### E-cadherin regulates epithelial cell cortex mechanics during apoptotic target engulfment

While we found that α/β-catenin was required for mechanical force transmission to mediate apoptotic cell engulfment, the formation of ‘epithelial arm’ protrusions that trigger apoptotic motility in vivo remained functional. In contrast, E-cadherin deficient embryos had a failure in both apoptotic target movement and clearance, suggesting a functional involvement of additional E-cadherin dependent molecular regulators.

Notably, a marked change in epithelial tissue architecture was evident in E-cadherin deficient embryos in regions with an increased apoptotic cell density, with epithelial cells showing a characteristic apical shrinkage associated with cell rounding, while a stretching of epithelial cells in adjacent tissue regions was observed (Supplementary Figure 6A,B). This heterogenous epithelial tissue remodelling did not occur in the absence of apoptotic cells (Supplementary Figure 3E,F), supporting that the change in tissue architecture was specifically depending on the contact with apoptotic cells in E-cadherin deficient epithelia.

Apical shrinkage and rounding of epithelial cells observed during apoptotic target encounter in E-cadherin deficient embryos was reminiscent of a change in cell mechanics and cortical contractility mediated by Myosin II activity^31^. To study if Myosin II is directly involved in phagocytic clearance, we first analysed the localization of Myosin II at the basal epithelial cell cortex during individual phagocytic uptake events. To exclusively visualize Myosin II-GFP dynamics at the basal epithelial cortex in contact with apoptotic cells, we performed transplantations of apoptotic targets from a donor embryo expressing a membrane marker (Lyn-tdTomato), in which cell death was induced by Bax overexpression, into a transgenic host embryo expressing Myosin II-GFP. In vivo high-resolution live cell imaging of interaction dynamics between epithelial and apoptotic cells revealed the transient accumulation of Myosin II at the local phagocyte-target contact site (Figure 6A). Myosin II localization followed a characteristic spatio-temporal pattern, with a progressive enrichment at the phagocytic cup during target engulfment until cup closure, and the rapid depletion of Myosin II upon target internalization and phagosome formation (Figure 6B; Supplementary Movie 20). No accumulation of Myosin II was observed at the basal epithelial cortex in the absence of apoptotic target contacts (Supplementary Figure 6C).

**Figure 6.**
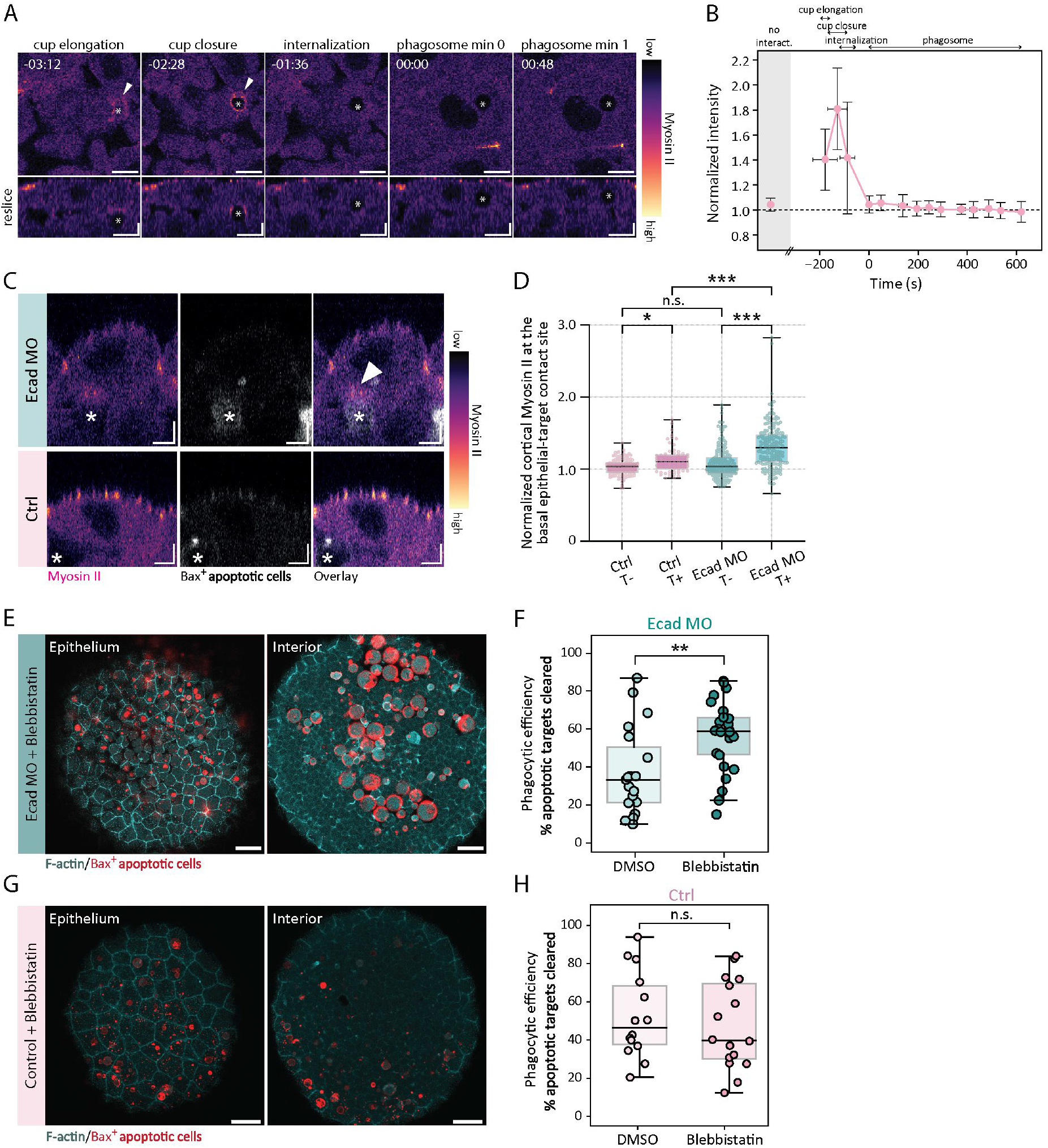
Spatio-temporal Myosin II dynamics at the phagocyte-target contact site and functional importance for efferocytosis. **A)** Dynamic localization of Myosin II (MRLC-GFP) at the basal epithelial surface (top) during engulfment (arrowheads) of an apoptotic cell (Bax+, asterisks) and transversal view (bottom). Time indicated in min:s. **B)** Quantification of the relative Myosin II intensity at the basal epithelial cortex before (grey area) and during (white area) phagocyte-target contact at indicated time points ( n=11, N=3); full engulfment at t=0. Data presented as mean+/-SD. **C)** Dual-colour fluorescence images (transversal view) of epithelial cells expressing MRLC-GFP (left) in contact with transplanted apoptotic targets (Bax+ cells, middle) and overlay (right) in control and Ecad MO embryos at shield stage (6 hpf). **D)** Quantification of relative cortical Myosin II levels at the epithelial-target interface in the absence (T-) or presence of transplanted apoptotic targets (T+) for control embryos (T-, n=130 measurements from 26 cells, 11 embryos/T+, n=64 measurements from 20 cells, 11 embryos) and Ecad MO embryos (T-, n=196 measurements from 47 cells, 23 embryos/T+, n=181 measurements from 49 cells, 23 embryos). N=3. Kruskall Wallis test with multiple comparisons, p=0.0121 (Ctrl T-/T+), p<10^-10^ (Ecad MO T-/T+), p=0.611 (Ctrl T-/Ecad MO T-), p=0.0002 (Ctrl T+/Ecad MO T+). **E-H)** Dual-colour fluorescence images (overlay) showing the location of apoptotic targets (Bax+ cells, red) in the epithelium (left) and embryo interior (right) in Ecad MO (E) versus control (G) embryos expressing Lifeact-GFP (cyan) at 75% epiboly (7.5 hpf) treated with Blebbistatin. Phagocytic efficiency in Ecad MO embryos (F) treated with DMSO (n=19 embryos) or Blebbistatin (n=26 embryos; unpaired t-test, p=0.0035) or control embryos (H) treated with DMSO (n=14 embryos) or Blebbistatin (n=16 embryos; Mann-Whitney test, p=0.7395). N=3. Embryos were obtained from Tg(actb2:myl12.1-EGFP) (A, C) and Tg(actb2:Lifeact-GFP) (E, G) lines. Large dots represent an embryo (C,F). N indicates the number of independent experiments. Box plot shows the median, the 1st and 3rd quartile and the min/max values (D) and the 1.5x inter-quartile range (F,H). All statistical tests are two-sided. Scale bars: x: 10 μm, y: 5 μm (A); x: 5 μm, y: 5 μm (C); 40 μm (E); 50 μm (G).

In contrast, the temporal localization dynamics of Myosin II were substantially altered in E-cadherin depleted embryos, with an excess and prolonged accumulation of Myosin II at the basal epithelial surface in contact with transplanted apoptotic targets (Bax+ cells) (Supplementary Figure 6E; Supplementary Movie 22), compared to a transient recruitment of Myosin II in control embryos (Supplementary Figure 6D; Supplementary Movie 21). This was concomitant with a pronounced local enrichment of Myosin II at the phagocyte-target contact site in E-cadherin deficient versus control embryos (Figure 6C,D). No accumulation of Myosin II was observed at the basal cortex of epithelial cells upon transplantation of live cells when in contact with the basal epithelial cell surface (Supplementary Figure 6F,G). These data support that E-cadherin is required to control the spatio-temporal dynamics of Myosin II at the local epithelial cell cortex in contact with apoptotic targets.

To directly validate that an increase in cell contractility limits apoptotic cell clearance in E-cadherin deficient tissues, we applied the Myosin II-specific inhibitor Blebbistatin. Treatment with Blebbistatin led to a typical reduction in epiboly movements during gastrulation, as described previously^32^, validating its effectiveness in vivo. Moreover, an increase in apical epithelial cell area in contact with apoptotic targets was observed, suggesting the contractile rounding phenotype in E-cadherin deficient condition was rescued (Supplementary Figure 7A). Importantly, we found that treatment with Blebbistatin was able to restore the phagocytic uptake defect in embryos injected with E-cadherin morpholino (Figure 6E,F). Consistently, the phagocytic clearance defect in DECMA-1 injected embryos was restored by Blebbistatin treatment (Supplementary Figure 7B,C). In contrast, wild type embryos treated with Blebbistatin had no significant difference in phagocytic clearance efficiency (Figure 6G,H). These observations support an important functional relation between E-cadherin and Myosin II activity that regulates epithelial efferocytosis.

Myosin II activity is controlled via reversible phosphorylation of its regulatory subunits^33^. To decrease Myosin II activity via a complementary genetic approach, we next overexpressed the Myosin Phosphatase (MP) complex based on the co-expression of the regulatory subunit (Mypt1) and two paralogs of the catalytic subunit Protein Phosphatase 1 (PP1βa and PP1βb)^34^. Overexpression of the MP complex in E-cadherin deficient embryos led to a significant increase in phagocytic clearance efficiency (Supplementary Figure 7D,E; Supplementary Movie 23), consistent with our results on the pharmacological inactivation of Myosin II activity via Blebbistatin.

Together, these results identify E-cadherin as a mechano-regulator of the phagocytic synapse, by controlling Myosin II activity at the epithelial cell cortex during the interaction and engulfment of apoptotic cells. An aberrant enrichment of Myosin II in E-cadherin deficient embryos is associated with a failure in apoptotic target clearance, supporting the spatio-temporal regulation of Myosin II activity at the basal epithelial surface is essential for proper apoptotic target clearance.

### E-cadherin signalling via p120 regulates Myosin II localization at the phagocytic synapse

We next aimed to address which signalling factors downstream of E-cadherin control Myosin II-dependent cell mechanics at the phagocyte-target contact site. P120-catenin has been implicated as a regulator of RhoGTPase activity^35,36^, and acts downstream of E-cadherin with a direct binding site to the cytoplasmic domain of E-cadherin at the juxtamembrane region^37^.

To support a direct functional link between p120-catenin and Myosin II localization at the phagocytic synapse, we next analysed the dynamics of p120-catenin at the basal epithelial surface upon apoptotic target encounter. We found that p120-catenin was dynamically recruited to the basal epithelial contact site during apoptotic cell interactions (Figure 7A; Supplementary Movie 24), where it transiently enriched during phagocytic cup progression and could also be detected on newly formed phagosomes (Figure 7A,B; Supplementary Movie 24). P120-catenin was not detected at the basal epithelial cell cortex in the absence of apoptotic cell interactions (Supplementary Figure 8A), and its recruitment failed in E-cadherin deficient conditions (Supplementary Figure 8B). These findings support that p120-catenin recruitment to the phagocytic synapse was directly mediated by E-cadherin (Figure 7C).

**Figure 7.**
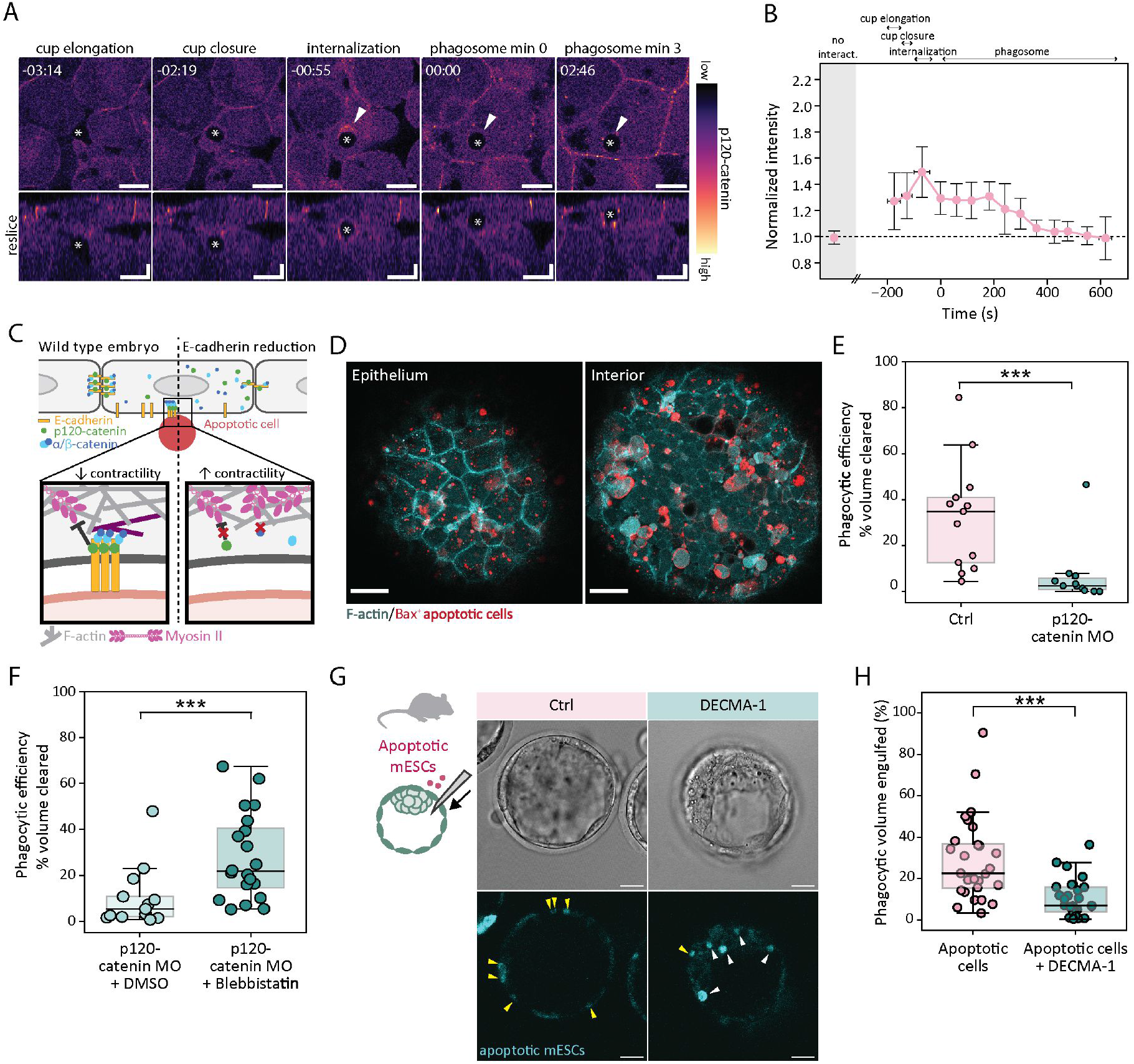
p120-catenin recruitment to the phagocytic synapse limits Myosin II levels for successful apoptotic cell clearance, and conservation of E-cadherin-dependent efferocytosis in mouse blastocysts. **A)** Fluorescence time lapse images showing GFP-p120-catenin recruitment in an epithelial cell during apoptotic cell engulfment (Bax+ cell, asterisks) at gastrula stage (5 hpf, top) and transversal view (bottom). Time indicated in min:s. **B)** Quantification of relative p120-catenin intensity at the basal epithelial surface before (grey area) and during phagocyte-target contact at indicated time points (white area; n=11, N=3). Data presented as mean+/-SD. **C)** The E-cadherin complex assembles at the basal epithelial phagocyte–apoptotic target contact site via recruitment of α-, β- and p120-catenin, coordinating successful target engulfment by locally restraining Myosin II activity and establishing a physical coupling to the actin cytoskeleton. Without catenin recruitment, actomyosin regulation is disrupted and engulfment compromised. **D)** Dual-colour fluorescence images (overlay) showing the location of apoptotic targets (Bax+ cells, red) in the epithelium (left) and the embryo interior (right) of a p120-catenin morphant embryo expressing Lifeact-GFP (cyan) at shield stage (6.5 hpf). **E)** Phagocytic efficiency (volume fraction) in control (Ctrl, n=13 embryos) and p120-catenin MO embryos (n=11 embryos). N=4. Mann-Whitney test, p=0.0005. **F)** Phagocytic efficiency (volume fraction) in p120-catenin MO embryos treated with DMSO (n=13 embryos) or 50 μM Blebbistatin (n=20 embryos) at 65% epiboly (7 hpf). N=3. Mann-Whitney test, p=0.0037. **G)** Representative images showing the localization of apoptotic mESCs (cyan, bottom) transplanted into the mouse blastocyst cavity (brightfield, top) in the absence (left) or presence of DECMA-1 antibody (right) 8 hours post transplantation (hpt) and **H**) quantification of the phagocytic efficiency in the absence (n=28 embryos) or presence of DECMA-1 (n=27 embryos) at 8 hpt. N=3. Welch two sample t-test; p=0.0001074. All embryos were obtained from the Tg(actb2:Lyn-tdTomato) (A) and Tg(actb2:Lifeact-GFP) (D) lines. Dots in (E,F,H) each represent an embryo. N indicates the number of independent experiments. Box plots show the median, the 1st and 3rd quartile and the 1.5x inter-quartile range. All used statistical tests are two-sided. Scale bars: x: 10 μm, y: 5 μm (A); 50 μm (D); 20 μm (G).

To assess its role in basal epithelial contact site mechanics, we depleted p120-catenin δ1, the most abundant p120-catenin isoform in early developmental stages^8,38^, via a previously validated morpholino^38^. Western Blot analysis confirmed a reduction of p120-catenin protein levels in morphant embryos (Supplementary Figure 8C). In contrast, junctional E-cadherin (Supplementary Figure 8D), and overall E-cadherin protein levels as assessed by Western Blot (Supplementary Figure 8E) were unaffected in p120-catenin deficient embryos. Notably, interfering with p120-catenin localization at the phagocytic synapse, by downregulating p120-catenin expression or its E-cadherin mediated recruitment, led to an aberrant enrichment of Myosin II at the basal epithelial cortex in contact with apoptotic targets (Supplementary Figure 8F, Supplementary Movie 25). The increased level of basal Myosin II was associated with a strong reduction in phagocytic clearance capacity (Figure 7D,E; Supplementary Movie 26), concomitant with a size reduction of residual phagosomes (Supplementary Figure 8G) and an accumulation of large apoptotic targets in the embryo interior (Supplementary Figure 8H). Moreover, p120-catenin depletion blocked the formation of epithelial ‘arms’ and thereby reduced apoptotic cell speed in vivo (Supplementary Figure 8I-L), leading to a restricted spatial dispersal of apoptotic targets.

Of relevance, treatment of p120-catenin deficient embryos with Blebbistatin, to inhibit Myosin II activation, was able to recover the phagocytic clearance defect (Figure 7F, Supplementary Figure 8M).

Interestingly, α-catenin depletion did not alter basal Myosin II levels (Supplementary Figure 9A; Supplementary Movie 27), and Blebbistatin treatment was not able to rescue the clearance defect in α-catenin depleted epithelia (Supplementary Figure 9B). These data validate the specificity of Myosin II inhibition and identify the E-cadherin/p120-catenin signalling complex as a critical regulator that limits Myosin II recruitment at the phagocytic synapse.

Together, these results support that the dysregulation of epithelial cell cortex mechanics at the phagocytic synapse causes a pronounced defect in apoptotic cell clearance by compromising functional protrusion formation and target engulfment.

### E-cadherin is required for apoptotic cell clearance in the mouse blastocyst

To study the functional role of E-cadherin in epithelial efferocytosis across species, we next assessed the physiological relevance of E-cadherin for apoptotic cell clearance in the early mouse embryo. We previously identified that the murine trophectoderm layer has a conserved phagocytic scavenging function mediating the removal of apoptotic cells in the mouse blastocyst^8^. To assess the functional role of E-cadherin in apoptotic cell clearance, we performed a transplantation of apoptotic mESCs into the mouse blastocyst cavity in the presence or absence of DECMA-1. Immunofluorescence imaging validated the localization of DECMA-1 at the basolateral domain of trophectodermal cells (Supplementary Figure 9C). Results showed that blastocyst mouse embryos co-injected with DECMA-1 had a compromised uptake of apoptotic cells, while apoptotic cells were engulfed by the trophectoderm layer in control embryos (Figure 7G,H).

These results support that E-cadherin is required for the clearance of apoptotic targets in mouse and zebrafish embryos, supporting a conserved function of E-cadherin in the phagocytic clearance of apoptotic cells by epithelia across species.

## Discussion

Epithelia perform various physiological functions in parallel to maintaining structural architecture and physical integrity. Epithelial phagocytosis represents a relevant physiological process in which a dynamic structural remodelling at the single epithelial cell level is required for target clearance, while tissue cohesion needs to be maintained. In this work, we show that epithelial cells can physically decouple phagocytic activities at their basal side from the apical domain, which ensures tissue architecture and stability. We identified that E-cadherin has a dual role in this process by providing both junctional stability and directly controlling the actomyosin cytoskeleton and mechanical force transmission at the phagocytic synapse. This extends the well-established role of E-cadherin at adherens junctions in epithelia and identifies a function of E-cadherin at the phagocyte-target contact site to mediate the clearance of apoptotic cells.

Several key molecular mechanisms underlying apoptotic cell recognition and uptake have been previously revealed and were shown to be conserved between professional and non-professional epithelial phagocytes. These include binding to phosphatidylserine (PS) on the surface of apoptotic cells ^39^ and the activation of Rac1 and PI3-kinase dependent signalling ^3,8,40^. Furthermore, the morphodynamic remodelling of the actin cytoskeleton and plasticity of professional phagocytes have been investigated^19,41^, including roles of mechano-transduction pathways in complement receptor (CR) mediated phagocytosis^42^. In contrast, the mechanisms underlying the morphodynamic remodelling and protrusion dynamics in non-professional epithelial cells have remained poorly understood.

Our data presented here support that the E-cadherin/catenin complex is repurposed at the basal surface in epithelial cells where it serves a critical role in the phagocytic clearance of apoptotic targets in vivo. We found that the E-cadherin/catenin complex forms de novo at the basal epithelial surface upon contact with apoptotic targets, with a fast recruitment of p120-catenin, α-catenin and β-catenin from the cytoplasm to the basal epithelial site in contact with apoptotic targets (Figure 7C). This dynamic assembly of the E-cadherin/catenin complex at the basal epithelial surface during interactions with apoptotic targets resembles the molecular events observed during the establishment of nascent cell-cell contacts, where E-cadherin engagement triggers the localized recruitment of catenins and actomyosin remodeling^29^. At nascent adhesion sites and mature junctions, E-cadherin assembles with catenins to form a signalling hub that provides both structural stability and a mechanical link to the actin cytoskeleton^16^. Similarly, we observed that the spatio-temporal dynamics of the E-cadherin/p120-catenin complex at the phagocytic synapse orchestrate localized actomyosin contractility, suggesting that epithelial cells co-opt the cadherin-catenin adhesion machinery for phagocytic clearance to regulate force transmission and actin/membrane remodelling to engulf apoptotic cells.

In previous work, we identified that apoptotic cell clearance by epithelial cells involves the formation of two types of dynamic protrusions at the basal epithelial surface: phagocytic cups, which mediate apoptotic target engulfment, and ‘epithelial arms,’ which generate mechanical pushing forces on apoptotic cells. The mechanical pushing forces generated by epithelial arms induce apoptotic target motility *in* vivo, which optimizes tissue clearance efficiency through a cooperative clearance activity among epithelial cells^8^. We further showed that these two types of epithelial protrusions mediate phagocytic clearance across different types of apoptotic targets, including spontaneous endogenous cell death, induced apoptosis or synthetic targets such as transplanted lipid aggregates that mimic essential apoptotic cell features but do not have intrinsic mobility^8^, supporting the validity of our experimental approach and the role of epithelial arms in apoptotic target movement in vivo. Synthetic apoptotic targets have further been widely used to study apoptotic cell clearance such as PS-coated beads or carboxylated beads^43,44^, supporting their experimental relevance. A key strength of our experimental model lies in its ability to enable the analysis of cell–cell interactions between phagocytes and apoptotic cells in vivo. Notably, we show here that the two types of epithelial protrusions are regulated in distinct manners by the E-cadherin/catenin complex: p120-catenin controls the local recruitment and temporal dynamics of Myosin II at the phagocytic contact site and its loss impairs both phagocytic cups and epithelial ‘arms’. In contrast, depletion of α-catenin or interfering with its actin binding site exclusively impairs the formation of phagocytic cups for target engulfment, while mechanical pushing through epithelial ‘arms’ remains functional. These findings suggest that α-catenin is essential for the transmission of actin polymerization forces along the target surface, which drives phagocytic cup progression. In contrast, pushing forces exerted by epithelial arms on the apoptotic target surface in the normal direction rely on perpendicular actin polymerization and form independently of α-catenin-mediated mechanical coupling to actin (Figure 5E). A similar critical role for force transmission has been identified for Integrin adhesion receptors in professional phagocytes^42^, supporting a wider involvement of adhesion receptors in mechanical force coupling during phagocytic clearance.

The mechanical state of the cell surface is influenced by Myosin II motor proteins that form bipolar thick filaments and bind to actin filaments to generate active forces which contract actin networks and thereby influences surface dynamics and protrusion formation^31,33^. Previous studies have addressed the mechano-signalling cross-talk between E-cadherin signalling and the actomyosin cell cortex at adherens junctions and its role in regulating cell mechanics in vitro and in vivo^45–47^. Myosin II activity is tightly regulated by phosphorylation of its regulatory subunits^33^. In this context, p120-catenin was shown to inhibit RhoA activity in epithelial cells via acting as a RhoGDI^48,49^. p120-catenin binds to the juxtamembrane region of the cytoplasmic tail of E-cadherin^37^ and can thereby function as a local modulator of RhoA and Myosin II activity. Furthermore, phosphorylation of p120-catenin through Src and Fyn tyrosine kinases can regulate the affinity of p120-catenin towards RhoA, possibly supporting a temporal control of RhoA activation upon receptor-ligand signalling activation^49^. Here we show that p120-catenin signalling directly controls the mechanics of the basal epithelial site in contact with apoptotic targets by regulating the spatio-temporal dynamics of Myosin II at the phagocytic synapse. Notably, downregulation of p120-catenin or interfering with its recruitment to the phagocytic contact site through the depletion of E-cadherin results in an aberrant recruitment of Myosin II, indicating a stiffening of the basal epithelial surface in contact with apoptotic cells. This excessive recruitment of Myosin II at the basal epithelial cortex hampers apoptotic cell clearance, and we show that specifically targeting Myosin II activity through pharmacological or genetic means is sufficient to recover apoptotic cell clearance.

The specific involvement of Myosin II in phagocytosis performed by professional phagocytes depends on the cell type and the biological context studied^50^. Higher target stiffness was shown to trigger an enrichment of Myosin II at the phagocytic synapse in macrophages^51^, and is required for late stages of phagocytosis, possibly involving phagocytic cup closure^52^. Myosin II and Myosin 1e/f were also implicated in having competing roles at the phagocytic cup^53^. Our findings support that the E-cadherin/catenin complex acts as a relevant mechano-regulator of phagocytic dynamics in non-professional epithelial phagocytes and directly controls the spatio-temporal recruitment of Myosin II at the phagocytic cup. Notably, our data show that an aberrant increase in Myosin II at the phagocytic synapse in epithelial cells compromises the phagocytic uptake of apoptotic cells, while the downregulation of Myosin II activity appears to have minor effects on the overall clearance efficiency. The local recruitment of the E-cadherin/catenin complex at the phagocytic contact site is thus functionally important to keep Myosin II activity in check and its de-regulation compromises the formation of functional protrusions required for apoptotic cell clearance. Together, our results establish a critical function of non-junctional E-cadherin/catenin complexes at the basal epithelial surface, controlling force transmission and the regulation of basal cortex mechanics to enable apoptotic cell clearance.

Functional relations between E-cadherin and the actomyosin cortex have been addressed in different biological processes in vivo. E-cadherin controls actomyosin flows at the cell and tissue level relevant in epithelial tissues^54,55^, establishes mechanical feedback during collective cell migration^56^ and provides adhesive couplings to generate traction forces for single cell migration^57^. Our findings show that E-cadherin also regulates the dynamics of the actomyosin cytoskeleton during epithelial phagocytosis. This is adding a mechanical function of E-cadherin in epithelia that connects tissue stability with dynamicity at the cellular level, associated to diverse physiological roles of epithelia during tissue development and homeostasis.

How the E-cadherin complex is activated at the phagocytic synapse remains to be addressed. From our data we can exclude that trans-binding is required for the de novo recruitment of the E-cadherin/catenin complex at the phagocyte-target contact site. Cis-clustering of E-cadherin molecules is a key step during the formation of nascent junctions^58^. Additionally, in non-junctional contexts like single cell migration^57^ or cytokinesis^59,60^, E-cadherin cis-interactions are sufficient to trigger the recruitment of catenins and can regulate actomyosin activity by means of the cadherin-catenin complex. E-cadherin thus fulfils important roles outside of cell-cell contacts and can modulate cellular mechanics in a cell autonomous manner. E-cadherin also has a large interactome, forming heterotypic cis-and trans-interactions^61^. The involvement of the EC1 domain based on our results further suggests interactions with ligands or receptor bridging molecules at the apoptotic cell surface. Other unconventional roles of E-cadherin have been identified in non-epithelial cells in the context of immune responses. These include the expression of E-cadherin on macrophages, which promotes the interaction and fusion of immune cells^62^ and modulates pro-inflammatory cell states^63^. E-cadherin has further been identified as a phagocytic receptor for the uptake of the pathogen *L. monocytogenes* by epithelial cells^64^. The involvement of cadherins in bacterial uptake^65^ and efferocytosis, as demonstrated in this study, together with their expression in close relatives of metazoans such as choanoflagellates, suggests an intriguing evolutionary role for E-cadherin in phagocytic prey uptake^66^.

During embryo development, the clearance of apoptotic cells by non-professional phagocytes further allows for the removal of aberrant and erroneous cells already from the earliest stages of development^8,67,68^ and enables the clearance of bacteria^65^, thereby establishing the earliest innate immune program in the developing embryo. Our work identifies that the E-cadherin/catenin complex forms a critical mechano-signalling pathway directly at the basal epithelial surface in contact with apoptotic targets that finely regulates basal epithelial cortex mechanics at the phagocytic synapse. Given the characteristic expression of E-cadherin in epithelial tissues and its high structural conservation, our findings suggest that E-cadherin acts as a mechano-regulatory receptor in apoptotic cell clearance in other epithelial tissue types and across species. Indeed, our data support that E-cadherin is required for apoptotic cell clearance in the early zebrafish and mouse blastocyst embryo, supporting its physiological relevance in lower vertebrates and mammals. The E-cadherin/catenin complex thereby establishes a critical mechano-signalling hub for regulating both physical tissue architecture and physiological tissue function, relevant to safeguard embryo development and to guarantee tissue homeostasis in the adult organism.

## Methods

### Zebrafish lines and maintenance

The following zebrafish lines were used in this study: AB wild type, Tg(actb2:Lifeact-GFP)^69^, Tg(actb2:Lyn-TdTomato)^70^, Tg(actb2:myl12.1-EGFP)^71^ and KI(mlanYFP)^xt17^cdh1-YFP embryos^72^ and Tg(Krt18:Gal4FF/UAS:Lifeact-GFP)^73^. Zebrafish were maintained at the aquatic facility of the Parc de Recerca Biomèdica de Barcelona (PRBB) following the standard procedures proved by the Institutional Animal Care and Use Ethic Committee (PRBB–IACUEC, CEEA-OH-PRBB:VRU-23-0014) according to national and European regulations. All the experiments were performed following the principles of the 3Rs. Eggs were kept in E3 medium (5 mM NaCl, 0.17 mM KCl, 0.33 mM CaCl_2_, 0.33 mM MgSO_4_) at 28°C or 31°C depending on the experiment. Embryos were staged based on morphological criteria and hours post fertilization (hpf)^74^ and stated in every figure legend. Sex of embryos is not reported since it is not yet determined during the developmental stage studied in this work.

### Plasmids cloning

#### GFP/mCherry p120 catenin cloning

To clone p120 *ctnnd1* (ENSDART00000106048) we used cDNA previously retrotranscribed (SuperScript™ III First-Strand Synthesis SuperMix for qRT-PCR, Invitrogen) from total mRNA isolated using 4 hpf zebrafish embryos (TripleXtractor direct RNA GK23.0100, Grispr). Phusion HF (Thermofisher F530S) was used to amplify the p120 sequence with the primers listed in Table 1. The Gibson Assembly System (in-house production) was used to introduce the p120 fragment into a pCS2 N-GFP or pCS2 N-mCherry vectors previously linearized with EcoRI/XbaI.

**Table 1.**
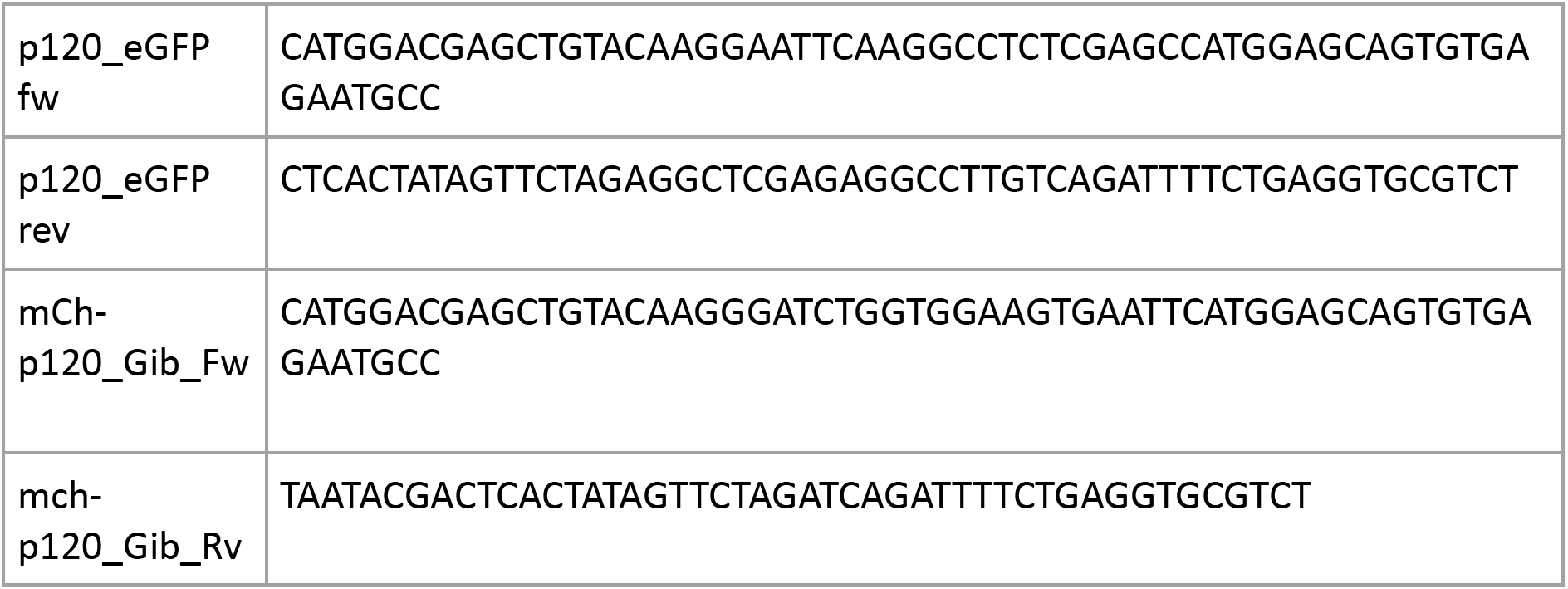
Primers used for cloning GFP/mCherry-p120 catenin.

#### α-cateninΔABS-mcherry cloning

To clone α-cateninΔABS-mcherry, zebrafish α-catenin (*ctnna*) lacking the actin binding site (aa 849-906) was amplified from pCS2 α-catenin-mCherry with the primers listed in Table 2. The Gibson Assembly System (in-house production) was used to introduce the α-catenin fragment into a pCS2 C-mCherry construct previously linearized with BamHI/EcoRI.

**Table 2.**
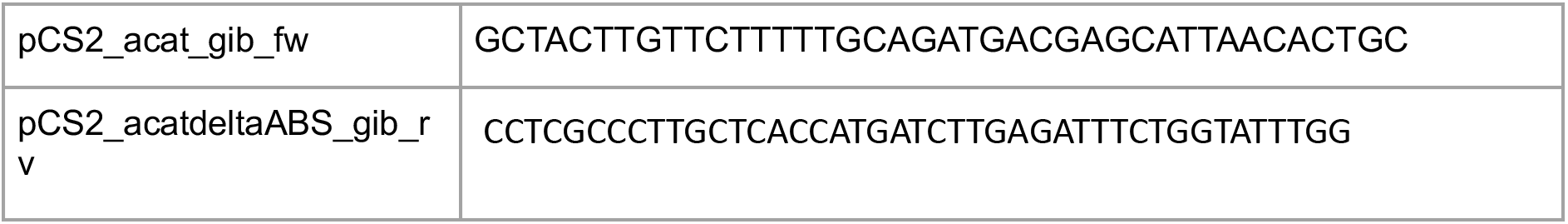
Primers used for cloning α-cateninΔABS-mCherry.

### mRNA synthesis and injections

mRNAs were synthetized from pCS2+ plasmids using the SP6 mMessage mMachine kit (Ambion, AM1340). Injections of mRNAs were performed as described in the following:

#### Induction of apoptosis

Apoptosis in a subpopulation of embryonic progenitor cells was induced by mosaic expression of *zbax* mRNA, as described previously. In brief, 2 pg of zbax mRNA was injected at 16-32–cell stage together with 100 pg of *lyn-tdTomato* mRNA to generate a mosaic expression of apoptotic cells (Bax+ cells) in embryos as indicated in each experiment. The used concentration of *zbax* mRNA led to the apoptosis of injected cells around dome stage (4.5 hpf) when embryos were selected based on the signal of Lyn-tdTomato expression and an increased Lifeact-GFP signal in apoptotic cells when the transgenic line Tg(actb2:Lifeact-GFP) was used. Embryos were then dechorionated and mounted for live in vivo imaging as described in detail (see ‘*Live cell in vivo microscopy*’).

#### Mypt complex injections

Tg(actb2:Lifeact-GFP) embryos were injected at 1-cell (zygote) stage into the yolk using a mix of 50 pg mypt mRNA, 50 pg pp1βa mRNA and 50 pg pp1βb mRNA^75^ in the presence or absence of 0.5 ng of E-cadherin morpholino (MO). Apoptosis of a subpopulation of embryonic progenitor cells was induced in the same embryos by mosaic expression of zbax mRNA, as described (see ‘Induction of apoptosis’).^768^

#### GFP/mCherry-p120-catenin, EGFP-β-catenin and α-catenin-mCherry injections

Tg(actb2:Lyn-tdTomato) line embryos were injected at 1-cell stage with 100 pg GFP-p120-catenin mRNA (control) or 100 pg GFP-p120-catenin mRNA and E-cadherin MO (EcadMO). Wildtype (AB) embryos were injected at 1-cell stage with 100 pg p120-catenin-mCherry mRNA, 100 pg EGFP-β-catenin mRNA or 100 pg α-catenin-mCherry mRNA.

#### α-catenin-ΔABSmCherry injections

Tg(actb2:Lifeact-GFP) embryos were injected at 1-cell (zygote) stage into the yolk with 400 pg α-catenin-ΔABSmCherry mRNA. Apoptosis of a subpopulation of embryonic progenitor cells was induced in the same embryos by mosaic expression of *zbax* mRNA, as described (see ‘*Induction of apoptosis*’).

### Use of morpholinos

Embryos were microinjected at 1-cell (zygote) stage into the yolk with translation-blocking morpholinos antisense oligonucleotides (Gene Tools). The following morpholinos (MOs) were used in this study:

E-cadherin MO^77^: ‘ATCCCACAGTTGTTACACAAGCCAT’.

p120-catenin MO^78^: ‘ACAATATGACCAGGGTCCAACATCC’.

α-catenin MO^79^: ‘CAAAATGGAGGGATGAGACTTTTAC’.

E-cadherin MO was injected at 0.5 ng per embryo (unless indicated otherwise), p120-catenin MO and α -catenin morpholino was injected at 4 ng per embryo.

### Pharmacological inhibitors and perturbation experiments

#### DECMA-1 injections

To inhibit E-cadherin binding activity in an acute manner, DECMA-1 anti-mouse-E-cadherin (BioXcell, CD324) was injected at a volume of 7-8 nl (4.67 mg/mL) into the blastula cup of embryos at dome stage (4.5 hpf) or at shield stage (6 hpf) as indicated. Injections were performed immediately prior to embryo mounting for live cell in vivo imaging. Embryos with drug treatments were first injected with DECMA-1 and then mounted and incubated for 2 hours prior to imaging. Control embryos were injected with a similar volume of DMEM medium (DMEM/F-12 with L-Glutamine and Hepes, Sigma). Embryos for immunostainings were injected with DECMA as described above at dome stage (4.5 hpf), incubated for 1h at 28°C and then fixed within the chorion in 4% PFA overnight at 4 °C.

#### Blebbistatin treatments

For inhibiting Myosin II activity, embryos with a mosaic subpopulation of apoptotic cells (see ‘*Induction of apoptosis*’) were selected and mounted (see ‘*Live cell in vivo Microscopy*’) in 1% low melting point agarose (UltraPure LMP Agarose, Invitrogen) prepared in Danieaús solution (58 mM NaCl, 0.7 mM KCl, 0.4 mM MgSO_4_, 0.6 mM Ca(NO_3_)_2_, 5 mM HEPES) containing 50 µM (S)-(-)-Blebbistatin (Tocris), or a similar volume of DMSO for control conditions. Samples were then incubated light-protected in Danieaús solution containing the same concentration of Blebbistatin or DMSO at 28°C for 3 h, corresponding to the time window during which most of the apoptotic cell clearance occurs. Finally, endpoint images were acquired.

### Preparation of lipid aggregates

POPS (1-palmitoyl-2-oleoyl-sn-glycero-3-phospho-L-serine, Avanti Polar Lipids), POPC (-palmitoyl-2-oleoyl-sn-glycero-3-phosphocholine, Avanti Polar Lipids) and Texas Red^TM^ DHPE (Texas Red^TM^ 1,2-Dihexadecanoyl-sn-Glycero-3-Phosphoethanolamine, Invitrogen) were dissolved in chloroform and mixed in a 80:19.9:0.1 molar ratio in a glass vial. The mix was dried under a gentle N_2_ flow and subsequently placed under high vacuum for 3 h. The lipid cake was suspended to a 20 mM lipid solution with PBS. The obtained aggregate suspension was washed two times with PBS by centrifugation of the solution for 1 min at 5000 g. Lipid aggregates were stored up to a few days at 4°C until usage.

### Zebrafish embryo transplantation experiments

#### Apoptotic cell transplantations

Apoptotic cell donor embryos were injected with 20 pg zbax and 100 pg lyn-tdTomato or lyn-Cerulean^80^ mRNA at 1-cell (zygote) stage (pCS-membrane-ceruleanFP was a gift from Sean Megason; Addgene plasmid #53749; http://n2t.net/addgene:53749; RRID:Addgene_53749). At sphere stage (4 hpf), embryos with a ubiquitous Lyn-tdTomato/Cerulean signal were selected as apoptotic cell donors. Apoptotic cells were transplanted to a localized region close to the epithelium of acceptor embryos at sphere to dome stage (4-4.5 hpf) using a Celltram Vario device (Eppendorf). An ES Blastocyst-pipette needle (Biomedical Instruments, VESbv-20-0-0-55) was used in combination with a custom-made transplantation agarose mold (Adaptive Science Tools). Donor cells were transferred into the blastula cup close to the epithelial layer and as far from the location where the needle was inserted as possible. Embryos were mounted 15 min after the transplantation to ensure wound healing prior to mounting.

The following zebrafish lines were used as donor and acceptor embryos in this study: Apoptotic cells with reduced E-cadherin expression were obtained from donor embryos from a Tg(actb2:Lifeact-GFP) line and embryos were injected with 1 ng of E-cadherin MO together with 20 pg zbax and 100 pg lyn-tdTomato mRNA into the yolk at 1-cell stage and transplanted into un-injected Tg(actb2:Lifeact-GFP) acceptor embryos.

To image the dynamic localization of E-cadherin during phagocytic cell clearance, wild type (AB) embryos were used as donor embryos for apoptotic cells and were injected at 1-cell stage with 20 pg zbax and 100 pg lyn-tdTomato mRNA. At sphere stage (4 hpf), embryos with a ubiquitous Lyn-tdTomato signal were selected as donors and apoptotic cells were transplanted into KI(mlanYFP)^xt17^cdh1-YFP acceptor embryos.

For basal Myosin II dynamic imaging and analysis, wildtype (AB) embryos were used as apoptotic cell donor embryos and injected at 1-cell stage with 20 pg zbax and 100 pg lyn-tdTomato or with 100 pg lyn-tdTomato mRNA for control experiments. At sphere stage (4 hpf), embryos with ubiquitous Lyn-tdTomato signal were selected as cell donors and cells were transplanted into E-cadherin MO injected or un-injected Tg(actb2:myl12.1-EGFP) embryos.

For p120-catenin, β-catenin and α-catenin localization experiments, wildtype (AB) embryos were used as apoptotic cell donor embryos and injected at 1-cell stage with 20 pg zbax and 100 pg lyn-Cerulean/lyn-tdTomato mRNA. At sphere stage (4 hpf), embryos with ubiquitous Lyn-Cerulean/tdTomato signal were selected as cell donors and cells were transplanted into GFP/mCherry-p120-catenin, GFP-β-catenin or α-catenin-mCherry injected acceptor embryos.

#### Transplantation of lipid aggregates

To mimic apoptotic cell clearance in vivo using synthetic targets, lipid aggregates were transplanted into control and E-cadherin morphant embryos at sphere stage (4 hpf) obtained from a Tg(actb2:Lifeact-GFP) line. For this purpose, lipid aggregates were first transferred into a transplantation agarose mold containing Danieau’s solution (without Ca^2+^ and Mg^2+^) in order to avoid further clustering of lipid aggregates and a clogging of the needle. Lipid aggregates were then transplanted into acceptor embryos using the same procedure as described for apoptotic cells (see section ‘*Apoptotic cell transplantations*’).

### Mouse blastocyst transplantation experiments

All protocols for mouse embryo experiments were performed in coordination with the CRG Tissue Engineering Facility and were approved by the Institutional Animal Care and Use Ethic Committee (PRBB–IACUEC), conforming to the guidelines of the European Community Directive and Spanish legislation for the experimental use of animals. Mouse lines were maintained and bred according to the standard procedures at the rodent facility of the Parc de Recerca Biomèdica de Barcelona (PRBB).

#### Apoptotic cell transplantation into mouse blastocyst embryos

Mouse blastocysts collection and cell transplantations were performed with the Tissue Engineering Unit at the CRG. The B6CBAF1/J and CD1 strains were used for transplantation of apoptotic cells. Embryos were isolated from B6CBAF1/J superovulated females or CD1 females (6-12 weeks old) mated with male mice (10-40 weeks old) of the same strain. Embryos were collected in M2 medium (Sigma, M1167) at 3.5 dpc and cultured in droplets of KSOM (Millipore, MR 106-D) covered with mineral oil (Sigma, M8410) at 37°C and 5% CO2.

For blastocyst transplantation experiments we used apoptotic mESCs (R1 H2B–mCherry/GFP–GPI^81^) obtained as a kind gift from the Hadjantonakis lab (Sloan Kettering Institute, US). Apoptotic mESCs were injected into the blastocyst cavity of 3.5 dpc embryos using an Olympus IX71 microscope and a Narishige MM-89 microinjector.

Prior to blastocyst transplantation, mESCs were maintained in KnockOut^TM^ DMEM medium supplemented with 10% ES qualified fetal bovine serum, 1x non-essential amino acids solutions, 1x Glutamax, 50 U/mL penicillin and 50 µg/mL streptomycin, 1x sodium pyruvate, 50 µM 2β-mercaptoethanol, and 1000 U/mL leukemia inhibitory factor (all from Life Technologies). Tissue culture-treated dishes (Falcon) were coated with 0.1% Gelatin (Sigma) for 30 min at 37 °C before seeding cells. Medium was changed every 24 h and cells were passaged every other day using 0.05% trypsin-EDTA (Life technologies) at ratios of 1:5 – 1:10. Cells were grown at 37 °C in an incubator with 5% CO2. For the induction of apoptosis, cells were plated in 24-well plate 5-7 days before blastocyst transplantation. The media was not changed during this period of time to induce apoptosis via nutrient-starvation. Prior to blastocyst transplantation, the supernatant of cell culture plates was evaluated at the microscope to identify wells with the most abundant level of cell death. The supernatant was collected and centrifuged at 300 g for 5 min. The pellet was either resuspended in 100 µL M2 media or 100 µL CD324 (E-cadherin) Monoclonal Antibody (DECMA-1). Cells were kept on ice until the blastocyst transplantation was performed.

### Fixation and immunostaining of mouse blastocyst embryos

Blastocysts were fixed 24 h post-transplantation for 10 min in 4% PFA. After 2 washes with PBS for 5 min, blastocysts were permeabilized with 0.5% Triton X-100 (Panreac) in PBS for 15 min, and then washed again 2 times in 0.1% Triton X-100 in PBS. To block unspecific binding, embryos were incubated in 3% BSA and 0.1% Triton X-100 in PBS for 45 min. Injected DECMA-1 antibody was detected via a secondary antibody Goat Anti-Rat IgG H&L - Alexa Fluor® 647 (ab150167, abcam, courtesy of Sdelci lab, CRG) that was incubated for 1 h at a concentration of 2µg/mL (1:1000 dilution from commercial stock). Embryos were co-stained with ∼0.165 µM Phalloidin-405 (Thermofisher, courtesy of Sebe lab, CRG). Embryos were washed 3 times for 5 min with 0.1% Triton X-100 in PBS. Fixed blastocysts were mounted in PBS drops on a bottom-glass 35 mm dish (FluoroDish, WPI, FD35-100), were covered with mineral oil (Sigma, M8410) and stored at 4°C in the dark until imaging.

### Fixation and immunostaining zebrafish blastula-gastrula embryos

For immunostaining, embryos obtained from the Tg(actb2:Lyn-tdTomato) and CRISPR/Cas9 knock-in line KI(mlanYFP)^xt17^cdh1-YFP line were used. Zebrafish embryos were staged and selected at 4 hpf, 6 hpf or 8 hpf and fixed within the chorion in 4% PFA overnight at 4 °C. Embryos were washed 3 times for 10 min with PBS-T (PBS + 0.1% Tween (Sigma)). Embryos were dechorionated and blocked in blocking buffer (PBS-T containing 10% goat serum and 2% BSA) for 4 hours at RT. For DECMA localization immunostaining, embryos were permeabilized with 2 washes in PDT (PBS-T + 0.3% Triton X-100 + 0.1% DMSO) for 30 min each. Embryos were incubated with primary antibodies diluted in blocking buffer overnight at 4°C (E-cadherin: 1:100, Biosciences 610181; PKC ζ (H-1): 1:200, Santa Cruz 1778). Embryos were washed 3 times for 10 min in PBS-T and blocked again in blocking buffer for 4 hours at RT. Embryos were incubated with secondary antibodies diluted in blocking buffer overnight at 4°C (anti-mouse Alexa-488, Invitrogen A-11001; anti-rat Alexa-647, abcam ab150167; phalloidin Alexa-405, Invitrogen A30104). Embryos were washed again 3 times for 10 min in PBS-T and kept in PBS until mounting.

### Western blot

To perform protein extraction, 150 embryos were deyolked at shield stage (6 hpf) per each condition following a previously described protocol^82^. Total cellular protein was extracted using RIPA digestion (Abcam ab156034), containing protease inhibitors (Roche 11836153001) and phosphatase inhibitors (Abcam ab201112). The protein amount was quantified using the BCA Protein Assay kit (Thermo Scientific 23227). A total amount of 40 µg of protein was separated by SDS-Page electrophoresis in a 10% acrylamide gel and transferred to a nitrocellulose membrane (Amersham 10600002). The membrane was incubated with primary antibodies for E-cadherin (Cdh1, 3:1000, BD Biosciences 610181), p120 (3:1000, Santa Cruz sc-23873) and Tubulin (1:1000, Sigma T9026) overnight at 4°C followed by 1 hour incubation at RT with a mouse secondary antibody (1:10000, Abcam ab205719). The SuperSignal kit (Thermo Scientific 34095) was used to develop the membranes for *cdh1* and p120-catenin and the ECL kit (Amersham RPN2232) was used for Tubulin. Pictures were acquired on the AI600 machine (Amersham) for E-cadherin and the iBright machine (Thermofisher) for p120-catenin.

### qPCR analysis

Control embryos and embryos injected with 0.5 ng or 4 ng of E-cadherin MO (3 independent replicates, each *n* = 30) were fixed with Trizol at sphere and shield stage (4 and 6 hpf) and placed in -20°C overnight. Samples were homogenized using a syringe and needle (BD Microlance, 0.4 mm x 19 mm 27G x ¾). RNA was prepared using the TripleXtractor Direct RNA kit (Grisp). 1 μg of total mRNA was used to synthesize cDNA (Xpert cDNA Synthesis Kit). 10 ng of cDNA was used to perform qPCR (Applied Biosystems 7900HT). The relative gene expression was normalized using *eef1a1* as a housekeeping gene and fold change was determined by applying the 2−ΔΔCt method.

### Live cell in vivo microscopy

#### Mounting of embryos for in vivo imaging

To perform 3D and 4D in vivo imaging, embryos were mounted in 1% low melting point agarose (UltraPure LMP Agarose, Invitrogen) in Danieaús solution on a 35 mm glass bottom dish with 10 or 14 mm inner diameter of the glass surface (MatTek) and covered with Danieaús solution to avoid sample drying. Selection of embryos for imaging was performed prior to imaging based on morphological criteria as the developmental stage, the absence of wounds in the epithelium and the preferential absence of Bax+ signal (based on Lyn-tdTomato expression) in the surface epithelium.

#### Microscopy

Embryos were imaged at Leica TCS SP5 CW-STED, SP8, SP8 FALCON STED or Zeiss LSM 980 with Airyscan 2 confocal microscopes. A laser excitation of 488 nm with a HyD, PMT, GaAsP-PMT or Airyscan GaAsP-PMT detector and 561 nm with a HyD, PMT or GaAsP-PMT detector were used. For KI(mlanYFP)^xt17^cdh1-YFP embryos, a laser excitation of 514 nm with PMT detectors was used. For Lyn-Cerulean apoptotic Bax+ targets, a laser excitation of 405 with a GaAsp-PMT detector was used. Live imaging of the clearance of apoptotic cells or lipid aggregates was performed with a HC PL APO CS2 20 x/0.75 NA immersion objective. High-resolution imaging of live and fixed embryos was performed with a HC PL APO CS2 63x/1.30 GLYC or HC PL APO CS2 40x/1.30 OIL objective. Live imaging of Myosin II dynamic accumulation in EcadMO, p120-catenin MO and control embryos, Myosin II transient localization and p120-catenin localization during the phagocytic uptake was performed with a LD LCI Plan-Apochromat 40x/1.2 Imm Korr DIC M27 objective (purchased through the Projecte BIST-AOM grant to ALMU) with silicon immersion media (Immersol Sil 406, Zeiss).

For live imaging of apoptotic clearance, z-stacks with a spacing of 1-2 μm between z-slices over a total depth of around 30 μm were acquired with a temporal resolution between 60 to 120 seconds depending on the experiment. For high-resolution imaging of the epithelial cell cortex and E-cadherin localization in live and fixed embryos, z-stacks with a spacing of 0.2 μm between z-slices were acquired. For Myosin II dynamic accumulation, Myosin II transient localization and p120-catenin localization during the phagocytic uptake z-stacks with a spacing of 1 μm between z-slices were acquired with a temporal resolution between 30 to 60 seconds depending on the experiment. For local basal Myosin II analysis, z-stacks with a spacing of 0.3 μm between z-slices were acquired.

Embryos were imaged at 28°C using an environmental incubator at the SP5 CW-STED, a H301-K stage top incubator (Okolab) with a UNO-T-H-CO2 controller (Okolab) at the SP8 and SP8 STED microscopes and an Ibidi Stage Top Incubator at the LSM980 Airyscan 2. Super-resolution imaging was performed on a ZEISS Elyra 7 microscope equipped with Zeiss lattice SIM and SIM² reconstruction. Images were acquired using a 63×/1.40 NA oil-immersion objective and an additional 1.6× tube lens. Excitation was achieved with a 488 nm laser line. All acquisitions were performed at a sample temperature of 28 °C. Z-step size was set according to Nyquist sampling (0.12 µm). Each SIM image was reconstructed from 13 raw phase images acquired with a square lattice pattern. Raw datasets were processed in ZEN Black software using the dual-iterative SIM² algorithm. In the first step, order combination, denoising, and frequency filtering were applied, generating a digital SIM point spread function (PSF). This PSF was then used for iterative deconvolution, improving both lateral and axial resolution compared to classical SIM reconstructions. Final reconstructions achieved an effective lateral resolution of ∼90 nm in XY.

#### Imaging of mouse blastocysts

For live imaging, injected blastocysts were kept in M2 medium droplets on a 35mm tissue culture dish with a No. 1.5 cover glass bottom (FluoroDish, WPI). Embryos were imaged with a HC PL APO CS2 20 x/0.75 NA glycerol objective on a Leica TCS SP8 confocal with 2x zoom. A laser excitation of 488 nm and/or 561 nm and PMT and HyD detectors were used. Samples were kept at 37°C and 5% CO2 using a H301-K stage top incubator (Okolab) with a UNO-T-H-CO2 controller (Okolab).

Fixed embryos were imaged with a HC PL APO CS2 20 x/0.75 NA glycerol objective on a Leica TCS SP8 confocal with 2.5x zoom. Laser excitation of 405 nm and 561 nm and PMT and HyD detectors were used.

### Image analysis

Image analysis was carried out with ImageJ^83^ and MATLAB (MathWorks, USA).

#### Analysis of phagosome size, target size and phagocytic efficiency

Identification and selection of apoptotic targets and phagosomes for analysis was based on their characteristic features: apoptotic targets were identified by their specific Lyn-tdTomato signal (Bax+ cells co-injected with the plasma membrane marker Lyn-tdTomato) together with morphological criteria as an irregular cell shape of apoptotic targets and/or an increased F-actin signal. Phagosomes were identified by their localization inside epithelial cells, their fluorescence signal (Lyn-tdTomato) and circular shape. The diameter was measured for phagosomes and the long and short axis was measured for apoptotic targets at their maximal cross-section using the *Line* tool. For the analysis of phagosome and target size, the diameter of all the phagosomes and targets above a threshold of 3 μm was included. All apoptotic targets in the field of view were included in the analysis, except for targets in contact with the yolk surface. For the analysis of phagocytic clearance in DECMA-1 injected embryos at shield stage (6 hpf), targets below a threshold of 8 μm were included to the analysis. The volume of apoptotic targets in the embryo interior was estimated assuming an ellipsoid geometry, while phagosome volume was estimated assuming a spherical geometry. The phagocytic efficiency was presented in two ways as indicated in the according data set: 1) the percentage of volume cleared by the embryonic epithelium from the total apoptotic volume present in the selected field of view in an embryo, or 2) as the percentage of the number of targets cleared by the epithelium calculated from the uptaken versus total number of available targets present in a selected field of view in an embryo. Embryos with an intact epithelium showing target engulfment and/or the presence of available apoptotic targets below the epithelium were included in the analysis.

#### Epithelial arm dynamics

Apoptotic target tracking was performed by using the *MTrackJ* plugin^84^. For control and E-cadherin MO embryos, a representative amount of motile apoptotic targets was tracked over time, with the track length limited by the duration of the time lapse imaging acquisition or targets moving out of field of view. The speed was calculated from the distance that targets moved in x-y-z direction between consecutive time points (time lag *t_lag_* = 90s). The maximum instantaneous speed for the given acquisition time was obtained from each track. For the visualization of the spatial target spreading in the embryo in vivo, the x- and y- coordinates of targets were obtained over a time period of 30 minutes from those tracks that exceeded at least once a max speed of 8.2 µm/min. The threshold was based on a previously derived mean target pushing speed^73^. Tracks were aligned and centred at the origin. For transplantation movies, apoptotic targets exceeding at least once a speed of 8.2 µm/min were tracked and included in the analysis. The speed was calculated from the distance that targets moved in x-y-z direction between consecutive time points (time lag *t_lag_* = 79-97s). The analysis of ‘high-speed’ targets was performed by calculating the mean speed of targets that were pushed between two events with high-speed (>8.2 µm/min) with a maximum duration of 10 min between these events. For the visualization of the spatial spreading of transplanted targets, the x- and y- coordinates were collected over a time period of 30 minutes from those tracks including at least two peaks of speed above 8.2 µm/min.

#### Frequency of phagocytic uptake and apoptotic cell movement events

A region of interest (ROI) of around 150 x 150 µm with available apoptotic targets was selected for each analysed embryo. All events of phagocytic target engulfment and movement were monitored over a duration of 120 min, identified by fast target movement accompanied by F-actin co-localization at the rear of the target for ‘arm’ pushing events, or by the appearance of phagosomes inside epithelial cells for phagocytic uptake events. The number of identified events within an interval of 30 min was divided by the total number of available targets in the selected ROI at the endpoint of each interval (30, 60, 90 and 120 min) and presented as the percentage of targets pushed or engulfed.

#### Analysis of junctional E-cadherin

To quantify the junctional intensity localization of KI E-cadherin-YFP from KI(mlanYFP)^xt17^cdh1-YFP embryos, a Gaussian blur filter (σ = 2) was applied and the intensity profiles normal to epithelial cell-cell junctions were obtained using the *Line* tool with a width of 5 pixels. The location of the junction was identified by the maximal intensity value and the intensity profiles were aligned at the centre according to the identified position of the junction.

To quantify junctional intensity of E-cadherin in immunofluorescence data, similar procedure was performed. Additionally, bleed through signal in this analysis was corrected by acquiring images of the Lyn-tdTomato signal with 488 nm excitation and subtraction of this signal from the E-cadherin signal detected with 488 nm excitation.

#### Analysis of apical epithelial cell area

The apical area of epithelial cells at different developmental stages was analysed in fixed embryos using the *Polygon Selection* tool. The apical cell area was measured as the 2D cross-sectional area of the apical epithelial surface. To analyse the apical area in epithelial cells in contact with apoptotic targets, epithelial cells from live embryos with a specific localization of Bax+ cells at their basal side were selected using the Lyn-tdTomato signal that was co-expressed in apoptotic targets. For apical cell shape analysis in the absence of apoptotic cells, epithelial cells were randomly selected for each field of view and their area measured or measured as indicated. For apical shape analysis of phagocytic and neighbouring non-phagocytic cells, phagocytic cells were chosen avoiding regions of high tissue stretching (due to epiboly) and non-phagocytic cells were selected among the neighbouring cells.

#### Analysis of local basal Myosin II localization and accumulation at phagocyte-target interface in epithelial cells

For the ‘local’ basal Myosin II analysis at the phagocyte target contact interface, epithelial cells in contact with individual apoptotic targets were first selected and a lateral reslice of the acquired z-stack was created. Basal Myosin II levels were quantified by manually identifying the basal surface of an epithelial cell and selecting the cortical region at the basal epithelial cell surface. Myosin II intensities were then analysed along a line of 10 pixels width for every 10 slices (with a spacing of 0.342 µm) using the *Line* tool, resulting in approximately 8 lines/cell. The average intensity values for each line were extracted to obtain a spatial intensity profile along the basal surface of epithelial cells. This intensity profile was normalized to the average cytoplasmic intensity that was obtained from 3 different slices for each cell. Normalized lines intensity profiles were obtained both at the local contact interface of an epithelial cell with an apoptotic target and at the basal epithelial site surrounding the contact site where no apoptotic targets were present. Data for contact sites with apoptotic (Bax+ cells) were then normalized to the median cortical signal in control embryos in the absence of targets and indicated as a ratio between cortical Myosin II values for apoptotic (Bax+ cells) and control (non-apoptotic) transplanted targets.

#### Analysis of Myosin II spatio-temporal localization and accumulation during phagocytic uptake in epithelial cells

Myosin II intensity levels were quantified by manually identifying uptake events and drawing a line of 1 pixel width along the cell membrane at the maximum cross-section of contact sites with targets using the *Segmented Line* tool. Specific timepoints of uptake (cup elongation, cup closure and internalization) were identified based on the position of the target compared to the cell. Phagosome uptake (time t=0) was identified as the stage at which the phagosome reached a final location in the epithelial cell and cell membrane returned to a resting stage. The average intensity values for each line were extracted to obtain a spatial intensity profile of Myosin II. The intensity profile at each timepoint was normalized to the average cytoplasmic intensity that was obtained from cytosol intensities measured around nuclear level with the *Oval selection* tool at different timepoints.

#### Analysis of p120-catenin spatio-temporal localization and accumulation during phagocytic uptake in epithelial cells

p120-catenin intensity levels were quantified by manually identifying uptake events and drawing a line of 1 pixel width along the cell membrane at the maximum cross-section of contact site with targets using the *Segmented Line* tool. Specific timepoints of uptake (cup elongation, cup closure and internalization) were identified based on the progression of cellular membrane around the target and position of the target compared to the cell. Phagosome uptake min 0 was identified as the stage at which the phagosome reaches a final location in the epithelial cell and cell membrane returns to resting stage. The average intensity values for each line were extracted to obtain a spatial intensity profile of p120-catenin. The intensity profile at each timepoint was normalized to the average cytoplasmic intensity that was obtained from cytosol intensities measured around nuclear level with the *Oval selection* tool at different timepoints.

#### Analysis of phagocytic efficiency of lipid aggregates

Quantification of phagocytic activity was performed using the *Multi-point* tool in FIJI. Large (> 20 µm) and small aggregates (< 3 µm) were excluded from the analysis. The phagocytic efficiency was calculated as the ratio of the number of lipid aggregates cleared by the embryonic epithelium and the total number of aggregates present in a given field of view.

#### Analysis of phagocytic efficiency of apoptotic cells in mouse blastocysts

Phagocytic efficiency was calculated as the ratio of the volume of phagosomes over the total volume of phagosomes and targets present in a given field of view. The volume of both apoptotic targets and phagosomes was estimated assuming a spherical geometry.

#### Optical flow analysis of Lifeact-GFP dynamics

Raw 3D SIM image stacks were processed in MATLAB (MathWorks) using the Bio-Formats MATLAB toolbox (bfmatlab). For each time-lapse dataset, maximum intensity projections were generated separately for the apical and basal regions of the EVL by selecting defined Z-slice ranges (∼0.5–6 µm depth). These projections were used as an input for flow analysis. Cytoskeletal dynamics were quantified using dense optical flow algorithms implemented in MATLAB. Specifically, the Farnebäck method (Computer Vision Toolbox) was applied to successive time frames to estimate per-pixel displacement vectors of the Lifeact-GFP signal. This yielded vector fields representing both direction and magnitude of actin flows. To reduce noise and highlight mesoscale patterns, raw vector fields were averaged over user-defined blocks (typically 16–32 pixels). Resulting flow fields were visualized as quiver plots overlaid on the corresponding SIM images (apical or basal projections) and as standalone magnitude heatmaps. For long time-lapse recordings, mean flow maps and temporal averages were calculated, providing robust measures of local actin flow dynamics across EVL cells.

### Statistical data analysis

The statistical tests used in each data set, *p*-values, the number of sample points n, and replicates N for each experiment are indicated in the figure legends. Statistical tests were performed using R and GraphPad Prism version 10.1.0. (for Windows and for Mac, GraphPad Software, Boston, Massachusetts, UAS, www.graphpad.com). All used statistical tests are two-sided. Data plots were generated using Python scripts (Python 3 reference manual^85^) and GraphPad Prism. SD indicates standard deviation and SEM the standard error of the mean. Box plots show the median, the 1st and 3rd quartile and the 1.5x inter-quartile range (IQR) unless indicated otherwise. p values are indicated as: * p<0.05, ** p<0.01, *** p<0.001. Source Data are provided as separate Source Data files.

## Data Availability

Source data are provided with this paper. Imaging raw data will be made available upon request.

## Code Availability

Custom-written Matlab code developed in this study is available under https://github.com/Stefan1980sol/ECAD.

## Acknowledgements

We thank L. Serrano lab for the PKC ζ (H-1) antibody, A. Sebé-Pedrós lab for the anti-mouse Alexa-488 secondary, S. Sdelci lab for the anti-rat Alexa-647 secondary, D. Weiser for Mypt, PP1βa and PP1βb plasmids, and C.P. Heisenberg for the α-catenin-mCherry and EGFP-β-catenin plasmids. This work was supported by the CRG Core Facilities for Advanced Light Microscopy, Tissue Engineering and Protein Technologies.

## Funding Statement

V.R. acknowledges financial support from the Ministerio de Ciencia y Innovacion through the Plan Nacional (PID2020-117011GB-I00), a research grant from HFSP (Ref.-No: RGY0079/2020), funding from the European Union’s Horizon Europe under the grant agreement No. BREAKDANCE_101072123 and funding (in part) by the Austrian Science Fund (FWF) 10.55776/PIN3977225. M.B. acknowledges funding from the Ministerio de Ciencia, Innovación y Universidades and Fondo Social Europeo (FSE) (PRE2020-092691). L.F.B acknowledges financial support from the “la Caixa” Foundation (ID100010434) with the fellowship code LCF/BQ/DI21/11860048.

## Author Contributions Statement

H-M.H., M.B. and L.F.B contributed to the research design, performed experiments and analysed data. S.J.-D performed Western Blots, qPCR analysis and plasmid cloning. F.P. performed plasmid cloning and contributed to mouse blastocyst experiments. S. Wieser performed SR live cell in vivo imaging and data analysis. L.C. contributed to morpholino interference experiments. C.V. provided scripts for data visualization. E.H. contributed to preliminary Blebbistatin experiments. S. Wijma contributed to catenin localization experiments. V.R. designed and performed research and supervised the project. V.R. wrote the manuscript with support of H-M.H., M.B and L.F.B.

## Competing Interests Statement

The authors declare no competing interests.

## Notes

### Competing Interest Statement

The authors have declared no competing interest.

### Summary of Updates

Revised formatting to match journal's requirements.

