## Supplementary Information File for "De novo E-cadherin/catenin complex formation controls basal epithelial mechanics and force transmission for apoptotic cell clearance"

[#] These authors contributed equally

#### Supplementary figures and legends

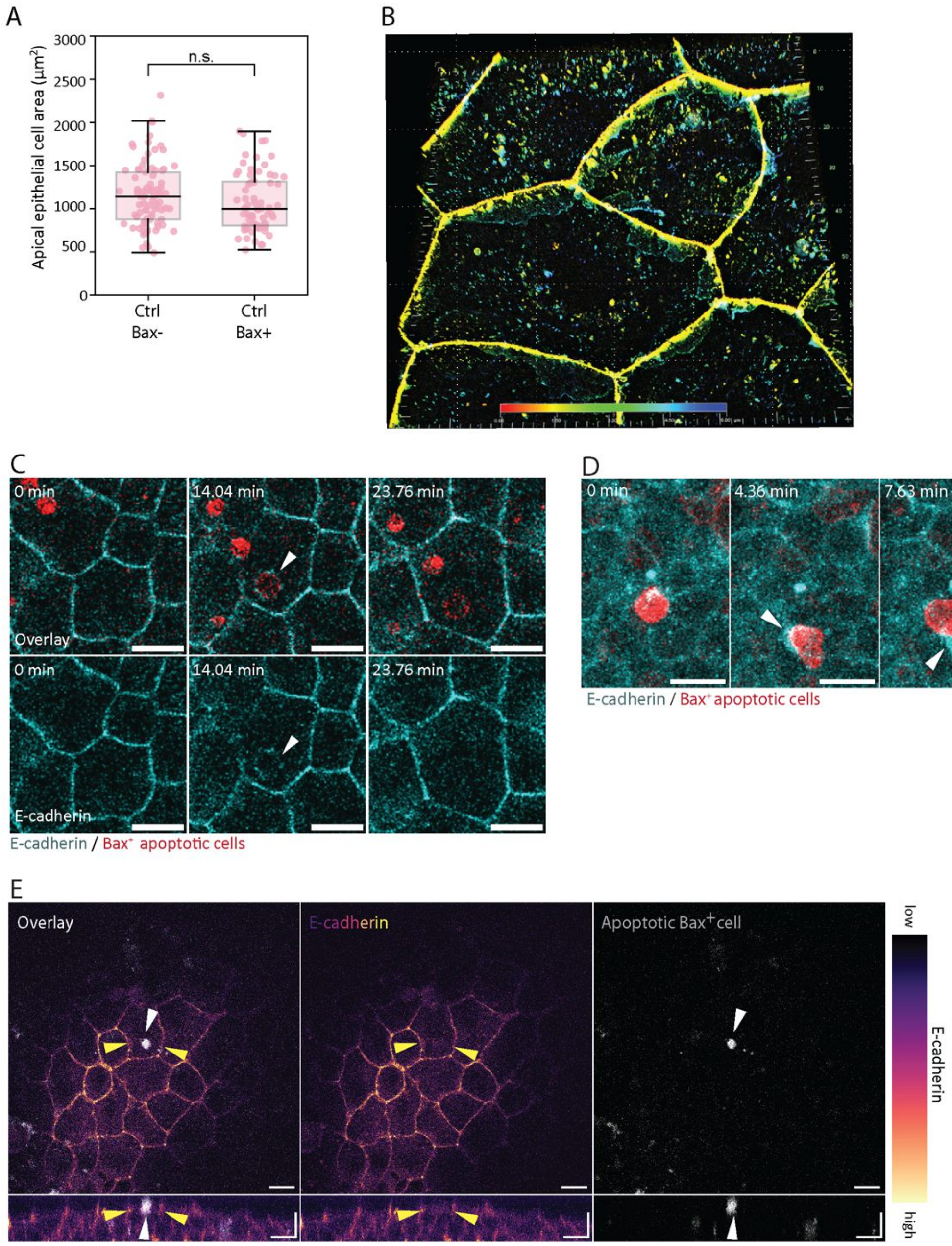


**Supplementary Figure 1. Basolateral localisation of E-cadherin in epithelial cells and E-cadherin localisation during apoptotic cell interaction and engulfment. A)** Measurement of apical epithelial cell area in the absence of apoptotic cell interactions (Bax-, *n*=85 cells) or upon contact of apoptotic cells overexpressing the proapoptotic factor Bax (Bax+, *n*=71 cells) in embryos at shield stage (6.5 hpf). *N*=3. Each dot represents an epithelial cell. Mann-Whitney test, *p*=0.1466. **B)** Super-resolution SIM imaging of E-cadherin-YFP at endogenous expression levels showing its baso-lateral localisation. The 3D projection is presented in false colours indicating the z-dimension from the apical (red, 0.0 µm) towards the basal (deep blue, -6 µm) surface of epithelial cells. Yellow (-1.5 µm) highlights the junctional E-cadherin ring present at cell-cell junctions. Towards the embryo interior, green (-3 µm) to blue colours highlight E-cadherin enrichment at the basal surface and ruffle-like membrane protrusions. **C)** Representative images showing the transient enrichment of E-cadherin (arrowhead) in a newly formed phagosome upon uptake of an apoptotic cell (top: Bax+ cells, red) in an embryo expressing E-cadherin-YFP (cyan) at endogenous levels. **D)** Representative dual-colour fluorescence images (overlay) showing the localisation of E-cadherin (arrowhead) at the interface between a transplanted apoptotic target (Bax+ cell, red) and an 'epithelial arm' protrusion in an embryo expressing a fluorescently tagged E-cadherin (cyan) at endogenous levels. **E)** Representative image (top) and transversal view (bottom) showing the localisation of E-cadherin-YFP, expressed at endogenous levels, at cell-cell junctions (yellow arrowheads) and at the basolateral side in epithelial cells that have engulfed transplanted apoptotic cells (Bax+ cells, grey, white arrowheads) at gastrula stage (7 hpf). Embryos were obtained from the Tg(actb2:Lifeact-GFP) line (B, C, D) and the KI(mlanYFP)xt17cdh1-YFP line (E). *N* indicates the number of independent experiments. Box plots show the median, the 1st and 3rd quartile and the 1.5x inter-quartile range. All used statistical tests are two-sided. Scale bars 10 µm (C, D); x: 20 µm, z: 20 µm (E).

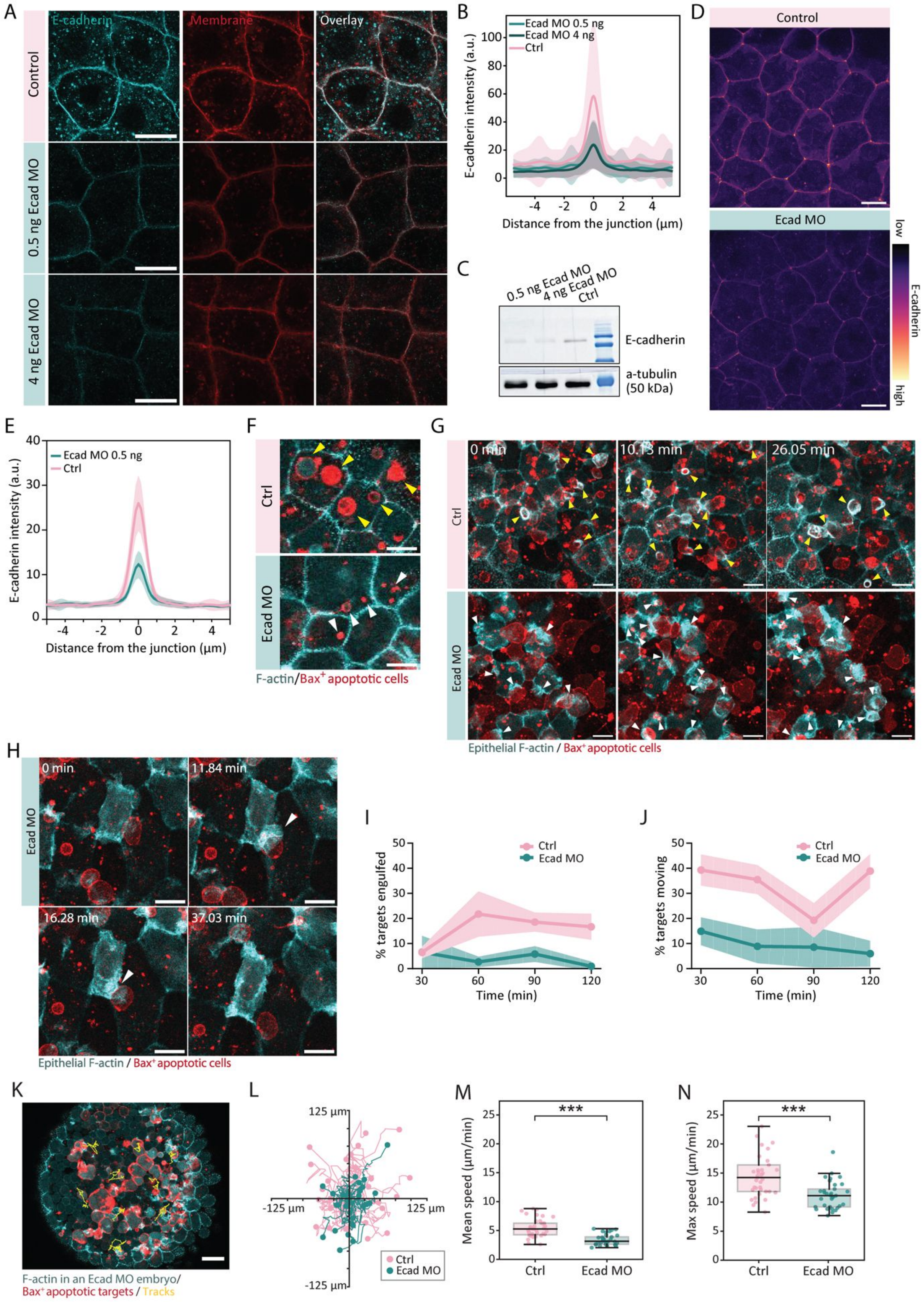


**Supplementary Figure 2. E-cadherin morpholino interference reduces E-cadherin levels in the embryonic epithelium and impairs apoptotic target engulfment and dispersal in vivo. A)** Representative immunofluorescence images showing the localisation of E-cadherin (cyan, left), the plasma membrane marker Lyn-tdTomato (red, middle) and merged images (right) in the epithelial tissue of control (top) and E-cadherin morphant (Ecad MO) embryos at shield stage (6 hpf) for two morpholino concentrations (0.5 ng, middle; 4 ng, bottom). **B)** Quantification of the E-cadherin signal intensity at epithelial cell junctions in control (pink, *n*= 62 junctions from 10 embryos, *N*=4), Ecad MO embryos injected with 0.5 ng morpholino (cyan; *n*=58 junctions from 12 embryos, *N*=4) and Ecad MO embryos injected with 4 ng morpholino (dark cyan; *n*=51 junctions from 10 embryos, *N*=3) at shield stage (6 hpf). Data lines and shaded areas represent the mean+/-SD. a.u. = arbitrary units. **C)** Western blot showing the reduction of E-cadherin in control (Ctrl) and Ecad MO embryos (injected with 0.5 ng and 4 ng morpholino) at shield stage (6 hpf). **D)** Representative images of epithelial cells at shield stage (6 hpf) in control and E-cadherin MO (0.5 ng) injected KI(mlanYFP)$^{xt17}$cdh1-YFP embryos. **E)** Quantification of the E-cadherin signal intensity at epithelial cell junctions in control (pink, *n*=15 junctions from 3 embryos, *N*=1) and Ecad MO embryos (cyan; *n*=15 junctions from 3 embryos, *N*=1) at shield stage (6 hpf). Data lines and shaded areas represent the mean+/-SD. a.u. = arbitrary units. **F)** Representative dual colour fluorescence images (overlay) of phagosomes (Bax+ cells, red) in epithelial cells of control (top) and Ecad MO (bottom) embryos expressing Lifeact-GFP (cyan) at shield stage (6 hpf). Arrowheads indicate individual phagosomes (yellow, Ctrl; white, Ecad MO). **G)** Time lapse dual-colour max z-projections images (Δz=4 µm; overlay) showing epithelial tissue F-actin dynamics in control (top) and E-cadherin MO injected (bottom) embryos expressing Lifeact-GFP (cyan) specifically in the epithelial layer under the krt18 promoter. Yellow arrowheads highlight successful phagocytic cups in control embryos, while white arrowheads point towards stalled phagocytic cups in E-cadherin MO embryos. **H)** Time lapse dual-colour max z-projections images (Δz=6 µm; overlay) of a single stalled phagocytic cup (white arrowhead) around an apoptotic cell (red) in an E-cadherin morphant embryo expressing Lifeact-GFP (cyan) specifically in the epithelial layer under the krt18 promoter. **I-J)** Percentage of targets within a selected field of view (approx. 150x150 µm) being engulfed by phagocytosis (I) and being pushed by epithelial ‘arm’ protrusions (J) in control (Ctrl) and E-cadherin morphant embryos (Ecad MO) from blastula to gastrula stage (4.5 - 6.5 hpf) (*n*=4 embryos (Ctrl), *n*=4 embryos (Ecad MO), *N*=2). Data show the mean+/-SEM. Multiple unpaired t-test, targets engulfed: *p*=0.9831 (30’), *p*=0.0813 (60’), *p*=0.0216 (90’), *p*=0.0206 (120’), targets pushed: *p*=0.0116 (30’), *p*=0.0075 (60’), *p*=0.2249 (90’), *p*=0.0040 (120’). **K)** Representative x/y-trajectories (yellow lines) obtained from 3D tracking of apoptotic cells (Bax+ cells, red) in an Ecad MO embryo expressing Lifeact-GFP (cyan) and visualized over a duration of 30 min. **L)** Tracks of individual apoptotic targets showing the path travelled over a period of 30 min in control (red, *n*=30 tracks from 6 embryos) and Ecad MO embryos (cyan, *n*=28 tracks from 5 embryos). Tracks were centred to the origin. Dots represent endpoint positions. **M)** Mean apoptotic cell target speed and **N)** maximum apoptotic target speed analysis in control (*n*=37 tracks from 6 embryos) and E-cadherin morphant (Ecad MO, *n*=33 tracks from 5 embryos) embryos from blastula stage (4.5 hpf) onwards. *N*=3. Data points represent individual tracks. Mann-Whitney test, $p<10^{-4}$. Embryos were obtained from the Tg(actb2:Lyn-tdTomato) line (A), the KI(mlanYFP)$^{xt17}$cdh1-YFP line (D), the Tg(actb2:Lifeact-GFP) line (F, K) and the Tg(Krt18:Gal4FF/UAS:Lifeact-GFP) line (G, H). *N* indicates the number of independent experiments. Box plots show the median, the 1st and 3rd quartile and the 1.5x inter-quartile range. All used statistical tests are two-sided. Scale bars: 10 µm (A); 20 µm (D, F, G,H); 50 µm (K).

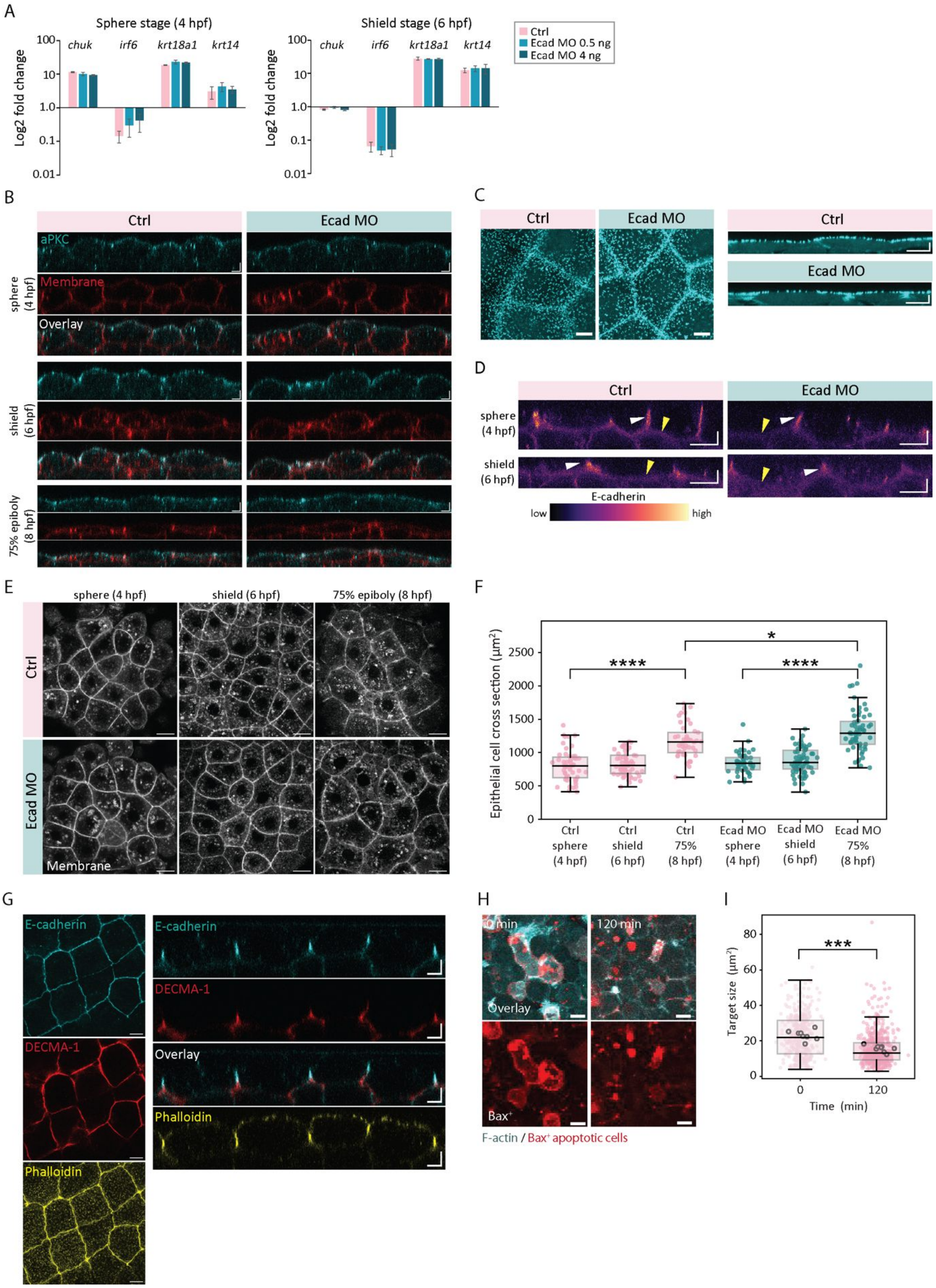


**Supplementary Figure 3. Reduction of E-cadherin expression or binding does not affect epithelial differentiation and architecture. A)** qPCR data showing the mRNA levels of epithelial specific markers (*chuk*, *irf6*, *krt18a1* and *krt4*) at sphere and shield stages (4/6 hpf) in control embryos (pink) and embryos injected with E-cadherin morpholino (0.5 ng, light blue; 4 ng dark blue). *N*=3. Data show the mean+/-SEM. **B)** Representative immunofluorescence images of the epithelium showing the localisation of the apical marker aPKC in control (left) and morphant (Ecad MO, right) embryos expressing a membrane marker (red, lyn-tdTomato) at sphere, shield and 75% epiboly stages (4/6/8 hpf). *N*=2. **C)** Representative maximum projection images of epithelial cells ($\Delta z$ = 1.6 µm) and transversal view (right) showing the localisation of actin at apical ridges and junctions in the epithelial tissue in control and E-cadherin morphant embryos expressing Lifeact-GFP (cyan) at shield stage (6 hpf). **D)** Maximum projection of re-sliced images ($\Delta z$ = 0.8 µm) of the epithelial layer showing the localisation of E-cadherin in control (left) and Ecad MO embryos (right) expressing fluorescently tagged E-cadherin at endogenous levels at sphere stage (4 hpf, top) and shield stage (6 hpf, bottom). White arrowheads indicate the localisation of E-cadherin at cell-cell epithelial junctions; yellow arrowheads indicate the basal localisation of E-cadherin. **E)** Representative immunofluorescence images of the epithelial tissue in control (top) and Ecad MO (bottom) embryos expressing a membrane marker (Lyn-tomato, grey) at sphere, shield and 75% epiboly stages (4/6/8 hpf). **F)** Quantification of the epithelial cell cross-sectional area at different developmental stages in control (Ctrl sphere stage, *n*=51 from 5 embryos, Ctrl shield stage, *n*=58 from 6 embryos, Ctrl 75% stage, *n*=45 from 6 embryos) and Ecad MO embryos (Ecad MO sphere stage, *n*=46 from 5 embryos, Ecad MO shield stage, *n*=69 from 7 embryos, Ecad MO 75% stage, *n*=60 from 8 embryos) in the absence of apoptotic cells. *N*=2. Each dot represents an epithelial cell. Mann-Whitney test, $p<10^{-4}$ (Ctrl sphere - Ctrl 75%), $p<10^{-4}$ (Ecad MO sphere - Ecad MO 75%), $p$=0.019 (Ctrl 75% - Ecad MO 75%). **G)** Representative transversal (right) and maximum projection (left) immunofluorescence images of the epithelium in embryos injected with DECMA-1 (red) at 50% epiboly (5.5 hpf), expressing E-cadherin-YFP (cyan) at endogenous levels and their overlay. Embryos were stained with phalloidin (yellow) to visualize the cell perimeter. DECMA-1 signal is present at the basolateral domain and does not interfere with cell-cell junctional integrity over the measurement duration of 1 hour. **H)** Representative dual-colour fluorescence images (overlay, left) showing a reduction of apoptotic cell size over time (Bax+ cells, red; right) in embryos with a ubiquitous expression of Lifeact-GFP (cyan) from blastula stage (4.5 hpf) to gastrula stage (6.5 hpf). **I)** Quantification of apoptotic cell size in embryos at blastula stage (4.5 hpf) and after 2 hours of cell clearance (shield stage, 6.5 hpf). These time points correspond to the stages of DECMA-1 injection (see also Figure 3F), supporting the clearance defect observed upon DECMA-1 injection occurs across a wide range of target sizes. Data were obtained from 8 embryos (*N*=3). Each dot represents an apoptotic target. Large dots represent the mean value for each embryo. Mann-Whitney test, $p<10^{-4}$. Embryos were obtained from the Tg(actb2:Lyn-tdTomato) line (B, E), Tg(actb2:Lifeact-GFP) line (C, H) and KI(mlanYFP)xt17cdh1-YFP line (D). *N* indicates the number of independent experiments. Box plots show the median, the 1st and 3rd quartile and the 1.5x inter-quartile range. All used statistical tests are two-sided. Scale bars: x: 10 µm, z: 10 µm (B, H); x: 10 µm, z: 5 µm (C, D); 20 µm (E); x: 5 µm, z: 5 µm (G, right); 10 µm (G, left).

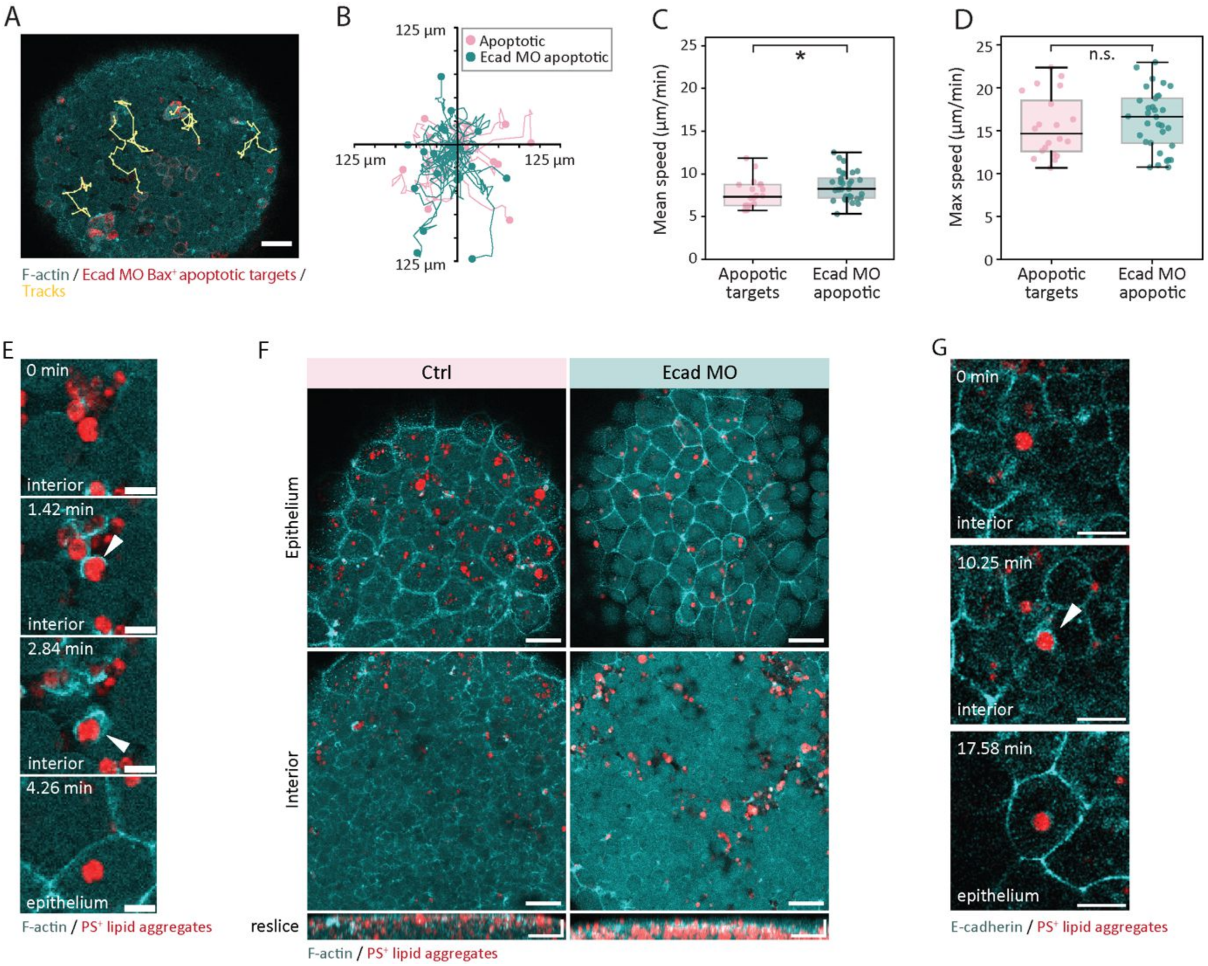


**Supplementary Figure 4. E-cadherin trans-binding is dispensable for apoptotic cell dispersal and uptake of synthetic targets. A)** Representative *x/y*-trajectories (yellow lines) obtained from 3D tracking of transplanted apoptotic cells (Bax+ cells, red) derived from a E-cadherin morphant (Ecad MO) donor embryo and transplanted into an embryo expressing Lifeact-GFP (cyan) and visualized over a duration of 30 min. **B)** Representative cell tracks showing the displacement of apoptotic cells transplanted from a control donor embryo (Ctrl, pink) and an E-cadherin morphant donor embryo (Ecad MO, cyan) into host embryos expressing Lifeact-GFP and visualized over a duration of 30 min. Trajectories were centred to the origin. Dots represent endpoints (*n*=14 tracks from 4 embryos (Ctrl), *n*=20 tracks from 6 embryos (Ecad MO). *N*=3. **C)** Mean speed and **D)** maximum speed analysis of apoptotic targets transplanted from control donor embryos (Ctrl, *n*=20 tracks from 4 embryos) and E-cadherin morphant (Ecad MO) donor embryos (*n*=33 tracks from 6 embryos) into host embryos expressing Lifeact-GFP. *N*=3. Each data point represents an individual track. Unpaired two-sided t-test; mean pushing speed: *p*=0.0302, max pushing speed: *p*=0.5003. **E)** Time lapse dual-colour images of F-actin localisation (arrowheads) in embryos expressing Lifeact-GFP (cyan) during the formation of a phagocytic cup engulfing a lipid aggregate (red containing Texas Red$^{TM}$-DHPE). **F)** Representative dual-colour fluorescence images (overlay) of the location of transplanted PS$^{+}$ lipid aggregates (red, containing Texas Red$^{TM}$-DHPE) in the epithelium (top) and embryo interior (bottom) of a control (Ctrl) embryo and E-cadherin morphant embryo (Ecad MO, 0.5 ng) expressing Lifeact-GFP (cyan) shown at 75% epiboly stage (8 hpf). **G)** Fluorescence time lapse dual-colour images of E-cadherin localisation (arrowheads) during the formation of a phagocytic cup engulfing a lipid aggregate (red, containing Texas Red$^{TM}$-DHPE) in an embryo expressing fluorescently tagged E-cadherin (cyan) at endogenous levels. Embryos were obtained from the Tg(actb2:Lifeact-GFP) line (A, E, F) and KI(mlanYFP)$^{xt17}$cdh1-YFP line (G). *N* indicates the number of independent experiments. Box plots show the median, the 1st and 3rd quartile and the 1.5x inter-quartile range. All used statistical tests are two-sided. Scale bars 50 µm (A); 10 µm (E); x: 40 µm, z: 20 µm (F); 20 µm (G).

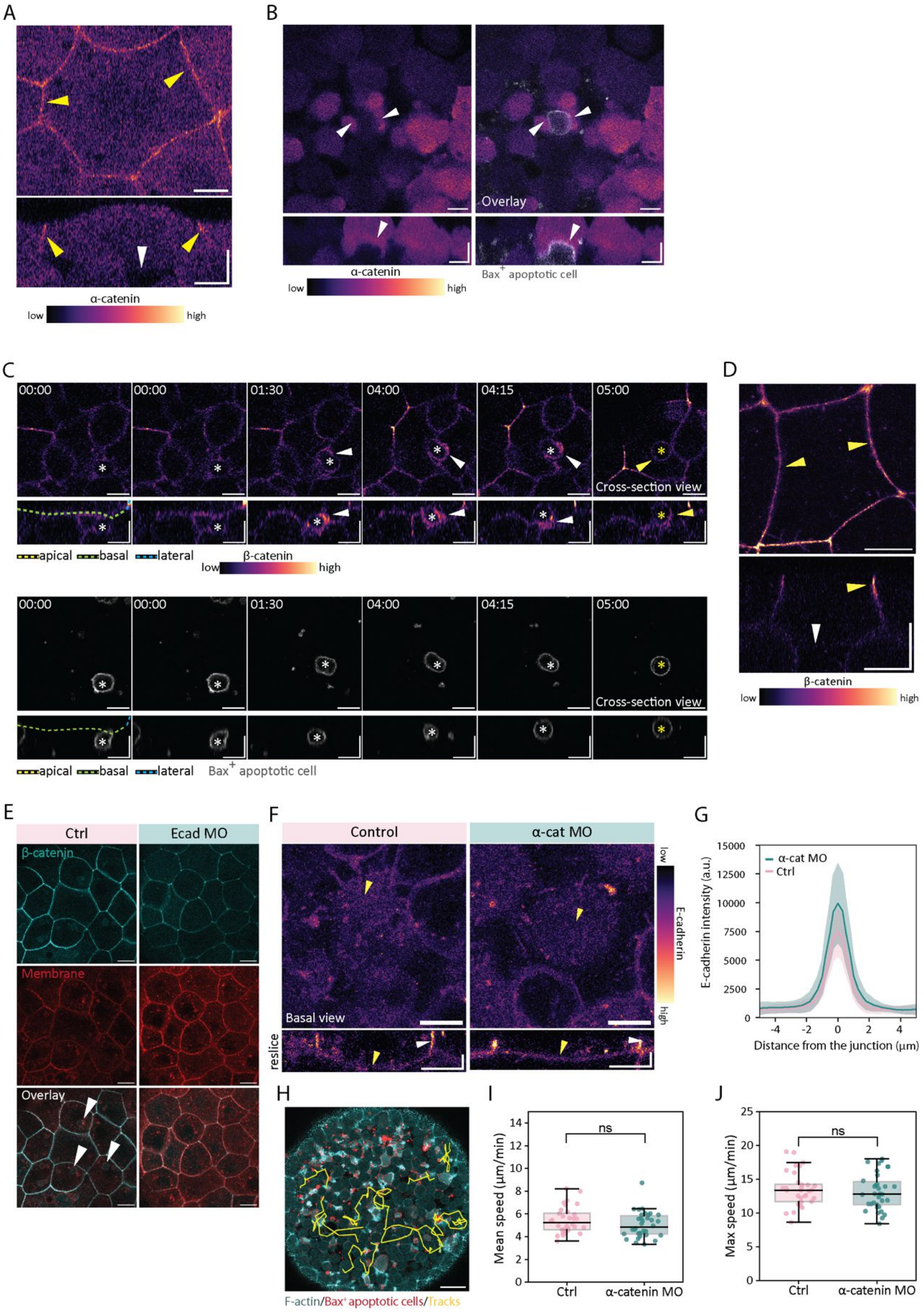


**Supplementary Figure 5. α- and β-catenin relocalise to the phagocytic synapse. Reduction of α-catenin expression levels does not affect apoptotic target pushing and dispersal in vivo. A)** Top view (top) and transversal view (bottom) of α-catenin-mCherry localisation in an epithelial cell in a wild type (WT) embryo at sphere stage (4 hpf), where it is present at cell-cell junctions (yellow arrowheads) but absent from the basal membrane (white arrowheads). **B**) Representative images showing the localisation of α-catenin-mCherry in epithelial cells in contact with transplanted apoptotic cells (Bax+ cells, grey) in E-cadherin morphant (Ecad MO) embryos at gastrula stage (5 hpf). Arrowheads indicate the absence of local enrichment of α-catenin-mCherry at the contact side with targets. **C)** Time lapse images showing the localisation of β-catenin (white arrowheads, top) during the uptake of a transplanted apoptotic cell target (Bax+ cells, grey, bottom) by epithelial cells in an embryo expressing EGFP-β-catenin. White arrowheads indicate β-catenin localisation during the uptake of an apoptotic cell, and yellow arrows show the localisation of β-catenin at the newly formed phagosome. Dashed lines outline the different surfaces of the phagocytic epithelial cell at time $t$=0. Time is indicated in min:s. **D**) Apical cross-section (top) and transversal view (bottom) of EGFP-β-catenin localisation in an epithelial cell in a WT embryo at sphere stage (4 hpf), where it is present at cell-cell junctions (yellow arrowheads) but absent from the basal membrane (white arrowheads). **E)** Representative images showing the expression of EGFP-β-catenin (cyan, top), the membrane marker Lyn-tdTomato (red, middle) and the overlay (right, bottom) in the epithelium of control embryos (left) and E-cadherin deficient embryos (right) at shield stage (6 hpf). EGFP-β-catenin localises at cell-cell junctions but does not accumulate in the nucleus even in cells that have engulfed endogenous apoptotic targets (white arrowheads in control embryos). **F)** Basal epithelial surface (top) and transversal view (bottom) showing E-cadherin localisation in an epithelial cell in control (left) and α-catenin morphant (right) embryos at shield stage (6 hpf). White arrowheads indicate the localisation of E-cadherin at cell-cell epithelial junctions; yellow arrowheads indicate the basal localisation of E-cadherin. **G)** Quantification of the E-cadherin signal intensity at epithelial cell junctions in control (pink, $n$=55 junctions from 6 embryos) and α-catenin morphant (α-cat MO) embryos (cyan; $n$=35 junctions from 5 embryos) at shield stage (6 hpf). $N$=2. Data lines and shaded areas represent the mean+/-SD. a.u. = arbitrary units. **H)** Representative x/y-trajectories (yellow lines) obtained from 3D tracking of apoptotic cells (Bax+ cells, red) in α-catenin morphant embryo expressing Lifeact-GFP (cyan) between (4.5 hpf) and shield (6.5 hpf) stage. **I)** Mean apoptotic cell speed and **J)** maximum apoptotic cell speed analysis in control ($n$=32 tracks from 3 embryos) and α-catenin morphant (α-catenin MO, $n$=34 tracks from 4 embryos) embryos from blastula stage (4.5 hpf) onwards. $N$=2. Data points represent individual tracks. Welch two-sample t-test, $p$=0.1466 (I), $p$=0.5868 (L). $N$ indicates the number of independent experiments. Box plots show the median, the 1st and 3rd quartile and the 1.5x inter-quartile range. All used statistical tests are two-sided. Embryos were obtained from the Tg(actb2:Lyn-tdTomato) line (E), the KI(mlanYFP)$^{xt17}$cdh1-YFP line (F) and the Tg(actb2:Lifeact-GFP) line (H). Scale bars x: 10 µm, z: 10 µm (A, B, C, D); 20 µm (E); x: 10 µm, z: 5 µm (F); 40 µm (H).

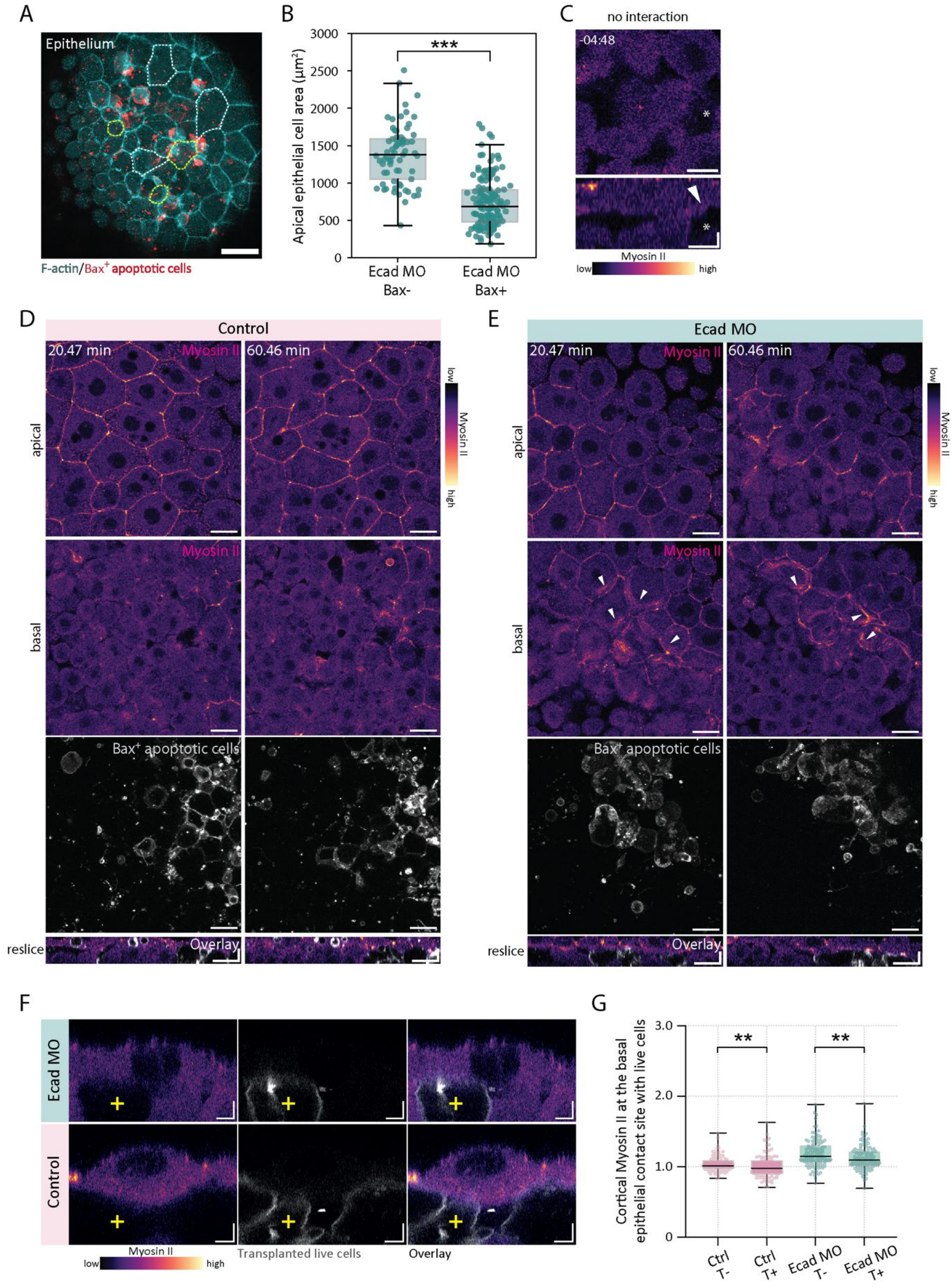


**Supplementary Figure 6. Dynamics and localisation of Myosin II in phagocytic epithelial cells at their basal surface depending on E-cadherin expression levels. A)** Representative dual-colour fluorescence image (overlay) showing changes in epithelial cell area upon contact with apoptotic targets (Bax+ cells, red) in an E-cadherin morphant embryo (Ecad MO) expressing Lifeact-GFP (cyan) at 75% epiboly stage (8 hpf). Yellow dashed lines indicate randomly selected epithelial cells in direct contact with apoptotic cells; white dashed lines indicate epithelial cells not in contact with apoptotic cells. **B)** Apical epithelial cell area in the absence (Bax-, *n*=69) or in direct contact with apoptotic cells (Bax+, *n*=149) in E-cadherin morphant (Ecad MO) embryos at shield stage (6.5 hpf). *N*=3. Each dot represents an epithelial cell. Mann-Whitney test, $p<10^{-4}$. **C)** Representative top view (top) and transversal view (bottom) showing the absence of Myosin II (MRLC-GFP) from the basal epithelial surface before contact (white arrowhead) with an apoptotic cells (Bax+, asterisk), which then will progress into a successful phagocytic uptake shown in Figure 6A at time t=0, representing the moment of apoptotic target internalization. **D-E)** Fluorescence time lapse images showing the localisation of MRLC-GFP at the apical domain (top) versus its accumulation on the basal epithelial surface (middle) at the contact sites with apoptotic cells (Bax+ cells, grey, bottom) in D) control and E) Ecad MO embryos at 30% epiboly (4.7 hpf, left) and 50% epiboly stage (5.5 hpf, right). Transversal view at the site of maximum target accumulation below the epithelial tissue (reslice). **F)** Representative fluorescence images in transversal view of epithelial cells expressing MRLC-GFP (left) in contact with transplanted live cells (middle; grey; yellow cross) and overlay (right) in control and E-cadherin morphant (Ecad MO) embryos at shield stage (controls) and 50% epiboly (6 hpf, Ecad MO). **G)** Quantification of cortical Myosin II at the phagocyte-target interface of epithelial cells in control embryos in the absence of targets (T-, *n*=91 measurements from 23 cells in 11 embryos) and in contact with transplanted control live cells (T+, *n*=95 measurements from 21 cells in 11 embryos) and Ecad MO embryos in the absence of targets (T-, *n*=121 measurements from 30 cells in 15 embryos) and in contact with transplanted control live cells (T+, *n*=123 measurements from 30 cells in 15 embryos). *N*=3. Mann-Whitney test, *p*=0.0064 (Ctrl), *p*=0.0062 (Ecad MO). *N* indicates the number of independent experiments. Box plot shows the median, the 1st and 3rd quartile and the 1.5x inter-quartile range (B) and the 1st and 3rd quartile and the min/max values (G). All used statistical tests are two-sided. Embryos were obtained from the Tg(actb2:Lifeact-GFP) line (A), and the Tg(actb2:myl12.1-EGFP) line (C, D, E, F). Scale bars: 50 µm (A); x: 10 µm, z: 5 µm (C); x: 20 µm (D, E, top views), x: 10 µm, z: 5 µm (D, E, reslices); x: 5 µm, z: 5 µm (F).

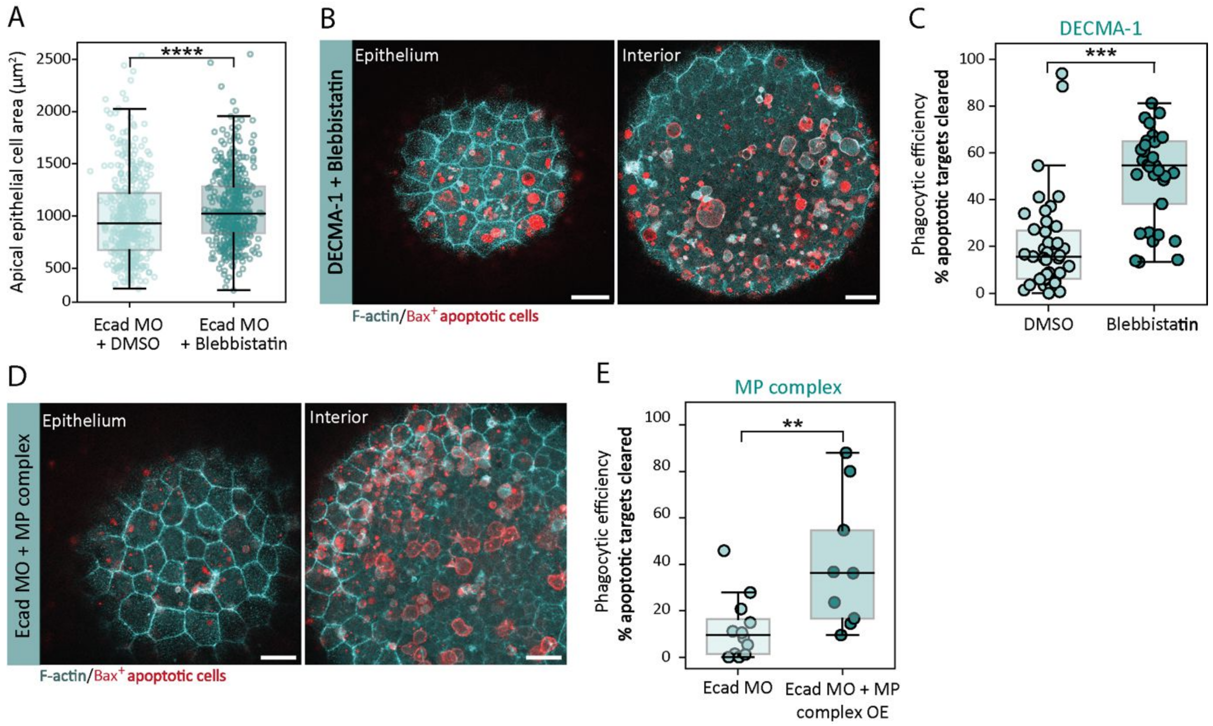


**Supplementary Figure 7. Phagocytic uptake of apoptotic cells is controlled by Myosin II activity. A)** Quantification of the apical area in epithelial cells in direct contact with apoptotic cells in Ecad MO embryos at 75% epiboly (7.5 hpf) incubated in DMSO (*n*=406 cells from 12 embryos) or in Blebbistatin (*n*=462 cells from 12 embryos). Mann-Whitney test, $p<10^{-4}$. *N*=3. **B**) Representative dual-colour fluorescence images (overlay) of the location of apoptotic targets (Bax+ cells, red) in the epithelium (left) and the embryo interior (right) at 70% epiboly stage (7 hpf) in DECMA-1 microinjected embryos incubated with Blebbistatin expressing Lifeact-GFP (cyan). **C)** Phagocytic efficiency derived as the percentage of apoptotic targets cleared by the epithelium in DECMA-1 injected embryos treated with DMSO (control, *n*=43) and treated with Blebbistatin (*n*=33). Unpaired two-sided t-test, $p<10^{-4}$. Each dot represents an embryo. *N*=3. **D)** Representative dual-colour fluorescence images (overlay) of the location of apoptotic targets (Bax+ cells, red) in the epithelium (left) and the embryo interior (right) at 70% epiboly (7 hpf) in Ecad MO embryos expressing Lifeact-GFP (cyan) and Myosin Phosphatase (MP) complex overexpression (OE). **E)** Phagocytic efficiency derived as the percentage of apoptotic targets cleared by the epithelium in Ecad MO (*n*=11) and Ecad MO + MP complex OE embryos (*n*=9). Data points represent individual embryos. *N*=3. Mann-Whitney test, *p*=0.0095. All embryos were obtained from the Tg(actb2:Lifeact-GFP) line. *N* indicates the number of independent experiments. Box plots show the median, the 1st and 3rd quartile and the 1.5x inter-quartile range. All used statistical tests are two-sided. Scale bars: 50 µm (B); 40 µm (D).

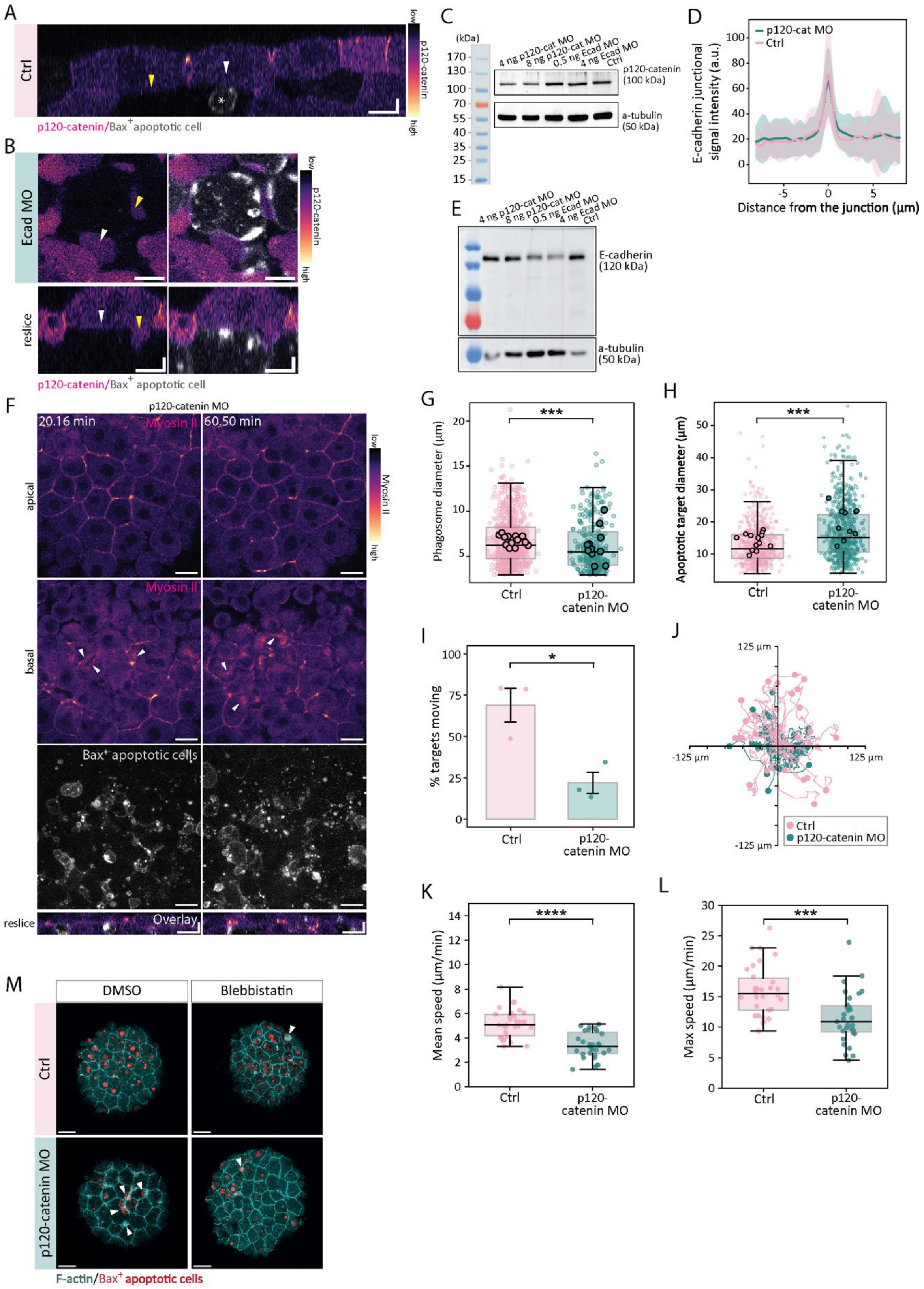


**Supplementary Figure 8. Reduction of p120-catenin does not affect E-cadherin expression and localisation but prevents phagocytic uptake and apoptotic target pushing**. **A)** Representative transversal views showing the absence of GFP-p120-catenin from the basal epithelial surface without apoptotic cell interactions (yellow arrowhead) and upon starting the interaction (white arrowhead)

with apoptotic cells (Bax+ cells, grey, asterisk). **B)** Representative dual-colour fluorescence images showing the localisation of GFP-p120-catenin (top left) at the phagocyte-target interface in E-cadherin morphant (Ecad MO) embryos, the overlay with apoptotic targets (Bax+ cells, grey, top right) and their transversal views (bottom). Arrowheads indicate the absence of a local enrichment of GFP-120-catenin at the basal epithelial surface (white arrowhead) and basal protrusion (yellow arrowhead) at contact sides with targets. **C)** Western blot showing the reduction of p120-catenin levels in p120-catenin MO embryos (4 ng and 8 ng) at shield stage (6hpf). p120-catenin levels remain similar in Ecad MO embryos (0.5 ng and 4 ng) and in control (Ctrl) embryos. **D)** Quantification of E-cadherin signal intensity at epithelial junctions in control (pink, *n*=75 junctions from 15 embryos, *N*=4) and p120-catenin MO embryos 4 ng (cyan; *n*=75 junctions from 15 embryos, *N*=4) at shield stage (6 hpf). Data lines and shaded areas represent the mean+/-SD. **E)** Western blot showing the levels of E-cadherin in p120-catenin MO embryos (4 ng and 8 ng), in Ecad MO embryos (0.5 ng and 4 ng) and in control (Ctrl) embryos at shield stage (6 hpf). a.u. = arbitrary units. **F)** Fluorescence time lapse images showing the localisation of Myosin II (MRLC-GFP) at the apical domain (top) versus its accumulation on the basal epithelial surface (middle) at the contact sites with apoptotic cells (bottom) in p120-catenin MO embryos at 30% epiboly (4.7 hpf, left) and 50% epiboly stage (5.5 hpf, right). Transversal view at the site of maximum target accumulation below the epithelial tissue (reslice). **G)** Phagosome diameter in control (Ctrl, *n*=702 phagosomes from 13 embryos) and p120-catenin MO embryos (*n*=300 phagosomes from 11 embryos) at shield stage (6.5 hpf). *N*=4. Each dot represents a phagosome. Mann-Whitney test, $p<10^{-4}$. **H)** Diameter of apoptotic targets in control (Ctrl, *n*=608 targets from 13 embryos) and p120-catenin MO embryos (*n*=720 targets from 11 embryos) at shield stage (6.5 hpf). *N*=4. Each dot represents a target. Mann-Whitney test, $p<10^{-4}$. **I)** Percentage of targets within a selected field of view (approx. 150x150 µm) being pushed by epithelial 'arm' protrusions in control (Ctrl) and p120-catenin morphant embryos (p120-catenin MO) from blastula to gastrula stage (4.5–5 hpf) (140 targets analysed from 3 embryos (Ctrl), 90 targets analysed from 3 embryos (p120-catenin MO), *N*=3). Data show the mean+/SEM. Each dot represents an embryo. Welch two-sample t-test *p*=0.024. **J)** Tracks of individual apoptotic targets showing the path travelled over a period of 30 min in control (pink, *n*=29 tracks from 3 embryos) and p120-catenin MO embryos (cyan, *n*=30 tracks from 3 embryos). Tracks were centred to the origin. Dots represent endpoint positions. **K)** Mean apoptotic cell target speed and **L)** maximum apoptotic target speed analysis in control (*n*= 0 tracks from 3 embryos) and p120-catenin morphant (p120-catenin MO, *n*= 1 tracks from 3 embryos) embryos from blastula stage (4.5 hpf) onwards. *N*=3. Data points represent individual tracks. Welch two-sample t-test $p<10^{-4}$ (K), *p*=0.00011 (L). **M)** Representative dual-colour fluorescence images of the location of apoptotic targets (Bax+ cells, red) in the epithelium at 7.5 hpf in control (top) and p120-catenin MO (4 ng, bottom) injected embryos expressing Lifeact-GFP (cyan) incubated with DMSO (control, left) or 50 µM Blebbistatin (right). White arrowheads indicate targets close to epithelial cells that are not cleared. *N* indicates the number of independent experiments. Embryos were obtained from the transgenic Tg(actb2:Lyn-tdTomato) (A,B), Tg(actb2:myl12.1-EGFP) (F) and Tg(actb2:Lifeact-GFP) (M) lines. Large dots in panels (G,H) represent the mean value for each embryo. *N* indicates the number of independent experiments. Box plots show the median, the 1st and 3rd quartile and the 1.5x inter-quartile range. All used statistical tests are two-sided. Scale bars: x: 10 µm, z: 5 µm (A, B); x: 20 µm, z: 10 µm (F); 50 µm (M).

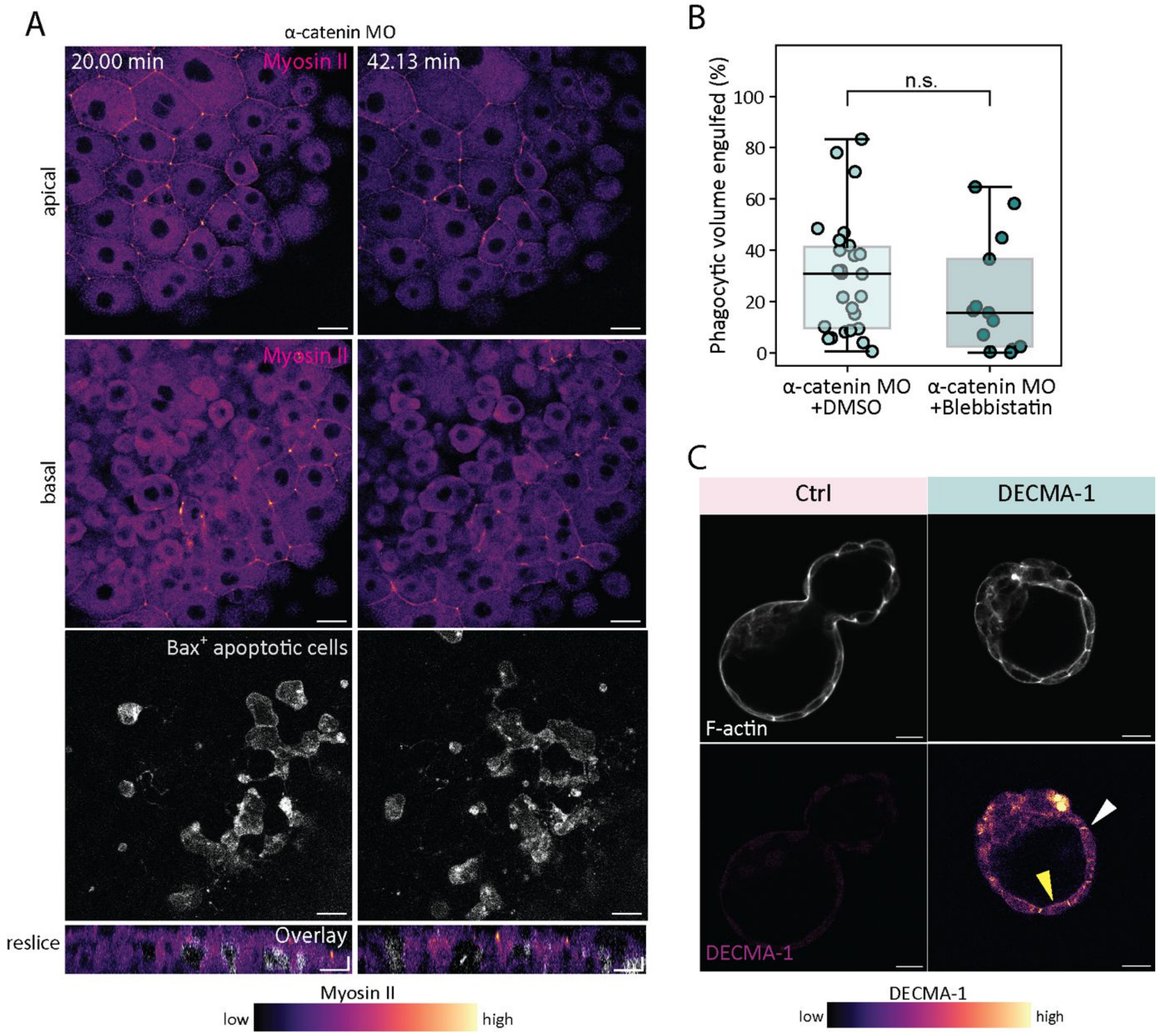


**Supplementary Figure 9. α-catenin does not control Myosin II dynamics at the phagocytic synapse. Validation of DECMA-1 localisation in mouse blastocysts. A)** Fluorescence time lapse images and transversal views showing the localisation of Myosin II (MRLC-GFP) at the apical (top) versus basal epithelial surface (middle) at the contact sites with apoptotic cells (bottom) in α-catenin MO embryos at 30% epiboly (4.7 hpf, left) and 50% epiboly stage (5.5 hpf, right). **B)** Quantification of the phagocytic efficiency for α-catenin deficient embryos treated with DMSO (α-catenin MO + DMSO, *n*=26) or Blebbistatin (α-catenin MO + Blebbistatin, *n*=13) at 7.5 hpf. *N*=3. Each dot represents an embryo. Welch two-sample t-test, *p*=0.2645. **C)** Representative images showing the basal (yellow arrowhead) and junctional (white arrowhead) localisation of an antibody directed against DECMA-1 (bottom) in fixed mouse blastocysts stained with phalloidin (top, grey) 24 hours after being injected with apoptotic mESCs in the absence (left) or presence of DECMA-1 antibody (right). *N* indicates the number of independent experiments. Box plots show the median, the 1st and 3rd quartile and the 1.5x inter-quartile range. All used statistical tests are two-sided. Scale bars: 40 µm (A), 20 µm (C).

### Supplementary Movie Legends

**Movie 1.** Time-lapse SIM imaging of the basal and apical side of the enveloping layer (EVL) in a zebrafish embryo at ~30% epiboly, expressing Lifeact-GFP to visualize filamentous actin. The movie shows dynamic actin flow patterns at the basal surface of epithelial cells versus a less dynamic actin organization at the apical cortex. Images were generated as maximum-intensity projections of z-stacks (~4.4 µm thickness, basal side) and (~0.4 µm thickness, apical side). Scale bars: 10 µm.

**Movie 2.** Phagocytic clearance in a control embryo (left) and E-cadherin morpholino injected embryo (Ecad MO, right) expressing Lifeact-GFP (cyan). Apoptotic targets (Bax+ cells) express a plasma membrane marker (red, Lyn-tdTomato). The phagocytic uptake of apoptotic targets by the embryonic surface epithelium (upper panel, *z* = 0 µm) and the phagocytic clearance dynamics in the embryo interior underneath the epithelium in a control (z = -14 µm) and an Ecad MO (*z* = -10 µm) embryo are shown. Embryos were imaged for 2 hours from 30% epiboly (4.7 hpf) to shield stage (6.5 hpf). Embryos were obtained from the Tg(actb2:Lifeact-GFP) line. Time is indicated in h:min:s:ms. Scale bar: 50 µm.

**Movie 3.** Phagocytic uptake events of apoptotic targets by single epithelial cells in an embryo expressing Lifeact-GFP (cyan). Apoptotic targets (Bax+ cells) express a plasma membrane marker (red, Lyn-tdTomato). Lines indicate the boundaries of the epithelial cell before (white) and just after (yellow) an apoptotic uptake event. The embryo was imaged from dome stage (4.5 hpf) to shield stage (6.5 hpf) and was obtained from the Tg(actb2:Lifeact-GFP) line. Time is indicated in min. Scale bar: 20 µm.

**Movie 4**. E-cadherin localization at two independent phagocytic cups (arrowheads) during the clearance of two apoptotic targets (red, Bax+ cells co-expressing the plasma membrane (PM) marker Lyn-tdTomato). The embryo was obtained from the CRISPR/Cas9 knock-in line KI(mlanYFP)$^{xt17}$cdh1-YFP expressing E-cadherin-YFP (cyan) at endogenous levels. Time is indicated in min. Scale bar: 20 µm.

**Movie 5.** Transient E-cadherin enrichment (arrowhead) in a newly formed phagosome upon uptake of an apoptotic cell (red, Bax+ cells co-expressing the plasma membrane (PM) marker Lyn-tdTomato). The embryo was obtained from the CRISPR/Cas9 knock-in line KI(mlanYFP)$^{xt17}$cdh1-YFP expressing E-cadherin-YFP (cyan) at endogenous levels. Time is indicated in min. Scale bar: 20 µm.

**Movie 6**. E-cadherin localization at an “epithelial arm” protrusion associated with an apoptotic target (red, Bax+ cells co-expressing the plasma membrane (PM) marker Lyn-tdTomato). The embryo was obtained from the CRISPR/Cas9 knock-in line KI(mlanYFP)$^{xt17}$cdh1-YFP expressing E-cadherin-YFP (cyan) at endogenous levels. Time is indicated in min. Scale bar: 20 µm.

**Movie 7.** Uptake of apoptotic targets and phagosome formation over time by single epithelial cells in a control (upper panel) and an E-cadherin morpholino (Ecad MO) injected embryo (lower panel) expressing Lifeact-GFP (cyan). Apoptotic targets (Bax+ cells) express a plasma membrane marker (red, Lyn-tdTomato). White lines indicate the boundaries of the same epithelial cell in control and Ecad MO embryos at the start and end point of the movie. White arrowheads indicate individual uptake events and the formation of a new phagosome. Embryos were imaged for 2 hours from 30% epiboly (4.7 hpf) to shield stage (6.5 hpf). Embryos were obtained from the Tg(actb2:Lifeact-GFP) line. Time is indicated in h:min:s:ms. Scale bar: 20 µm.

**Movie 8**. Epithelial tissue and single cell F-actin dynamics (cyan) in the presence of apoptotic cells (red, Bax+ cells co-expressing the plasma membrane marker Lyn-tdTomato) in control (left) and E-cadherin MO injected (right) embryos. Embryos were obtained from the Tg(Krt18:Gal4FF/UAS:Lifeact-GFP) line, which expressed Lifeact-GFP exclusively in the epithelial tissue. Time is indicated in h:min:s. Scale bar: 20 µm.

**Movie 9.** In vivo tracking (yellow lines) of apoptotic targets (red, Bax+ cells co-expressing the plasma membrane marker Lyn-tdTomato) in a control embryo (left) and an E-cadherin deficient embryo (Ecad MO, right) obtained from the Tg(actb2:Lifeact-GFP) line. Time is indicated in h:min:s. Scale bar: 50 µm.

**Movie 10.** Phagocytic clearance in a control (left) and a DECMA-1 injected embryos (right) expressing Lifeact-GFP (cyan). Apoptotic targets (Bax+ cells) express a plasma membrane marker (red, Lyn-tdTomato). The phagocytic uptake of apoptotic targets by the embryo epithelium (upper panel, $z$ = 0 µm) and phagocytic clearance dynamics in the embryo interior underneath the epithelium in a control ($z$ = -8 µm) and a DECMA-1 injected embryo ($z$ = -14 µm) are shown. Embryos were imaged for 2 hours from 30% epiboly (4.7 hpf) to shield stage (6.5 hpf). Embryos were obtained from the Tg(actb2:Lifeact-GFP) line. Time is indicated in h:min:s:ms. Scale bar: 50 µm.

**Movie 11.** Phagocytic clearance in control (left) and DECMA-1 injected embryos (right) expressing Lifeact-GFP after apoptotic cell fragmentation occurred at 50% epiboly stage (5.5 hpf). Apoptotic targets express a plasma membrane marker (red, Lyn-tdTomato). The upper panel shows a maximum-intensity projection of the embryonic epithelium and the location of phagosomes before DECMA-1 injection and a small amount of newly formed phagosomes after 2 hours of imaging (both indicated by white arrowheads). The lower panel shows the embryo interior ($z$ = -10 µm) underneath the epithelium, supporting the availability of small apoptotic targets (yellow arrowheads) in the embryo interior. Embryos were imaged for 2 hours from 30% epiboly (4.7 hpf) to shield stage (6.5 hpf). Embryos were obtained from the Tg(actb2:Lifeact-GFP) line. Time is indicated in h:min:s:ms. Scale bar: 50 µm.

**Movie 12.** Phagocytic clearance of transplanted apoptotic cells from a control donor embryo (left) or a donor embryo injected with E-cadherin morpholino (Ecad MO, right) into a host embryo expressing Lifeact-GFP (cyan). Apoptotic targets (Bax+ cells) express a plasma membrane marker (red, Lyn-tdTomato). The upper panel shows the phagocytic uptake of apoptotic targets by the embryo epithelium. The lower panel shows the phagocytic clearance dynamics in the embryo interior for control apoptotic cells ($z$ = -16 µm) and apoptotic cells obtained from an Ecad MO donor embryo ($z$ = -12 µm). Embryos were imaged for 2 hours from 30% epiboly (4.7 hpf) to shield stage (6.5 hpf). Embryos were obtained from the Tg(actb2:Lifeact-GFP) line. Time is indicated in h:min:s:ms. Scale bar: 50 µm.

**Movie 13.** Phagocytic clearance of transplanted PS+ lipid aggregates (red, synthetic apoptotic targets) in a control embryo (left) and an E-cadherin morphant embryo (Ecad MO, right) expressing Lifeact-GFP (cyan). The phagocytic uptake of synthetic apoptotic targets by the embryonic epithelium (upper panel) and the phagocytic clearance dynamics in the embryo interior (lower panel) are shown. Embryos were imaged over a duration of 3 hours from 30% epiboly (4.7 hpf) to 75% epiboly stage (8 hpf). Embryos were obtained from the Tg(actb2:Lifeact-GFP) line. Time is indicated in h:min:s. Scale bar: 40 µm.

**Movie 14.** E-cadherin localization at a phagocytic cup (arrowheads) during the clearance of a lipid aggregate (red) from the embryo interior (bottom) to the epithelium (top). Embryo was obtained from the CRISPR/Cas9 knock-in line KI(mlanYFP)xt17cdh1-YFP expressing E-cadherin-YFP (cyan) at endogenous levels. Time is indicated in min. Scale bar: 20 µm.

**Movie 15.** Localization of α-catenin-mCherry (white arrowheads, left) during the uptake of an apoptotic cell target (Bax+ cells co-expressing the PM marker Lyn-Cerulean, asterisks, right) by epithelial cells. The upper panel shows a top view and the bottom panel shows a transversal view of epithelial cells. White asterisks indicate the apoptotic cell targets within the embryo interior, while the yellow asterisk mark the engulfed phagosome. Time is indicated in h:min:s. Scale bar 10 µm.

**Movie 16.** Localization of β-catenin-GFP (white arrowheads, left) during the uptake of an apoptotic cell target (Bax+ cells co-expressing the PM marker Lyn-tdTomato, asterisks, right) by epithelial cells. The upper panel shows a top view and the bottom panel shows a transversal view of epithelial cells. White asterisks indicate the apoptotic cell target within the embryo interior, while yellow asterisk marks the engulfed phagosome. Time is indicated in h:min:s. Scale bar 10 µm.

**Movie 17.** Phagocytic clearance of apoptotic targets (Bax+ cells co-expressing the PM marker Lyn-tdTomato, red) in control (left) and α-catenin morphant (MO, right) embryos expressing Lifeact-GFP (cyan). The upper panel shows the phagocytic uptake of apoptotic targets by the embryonic epithelium. The lower panel shows the phagocytic clearance dynamics within the embryo interior. Embryos were imaged for 2 hours from 30% epiboly (4.7 hpf) to shield stage (6.5 hpf) and were obtained from the Tg(actb2:Lifeact-GFP) line. Time is indicated in h:min:s. Scale bar 40 µm.

**Movie 18.** In vivo tracking (yellow lines) of apoptotic targets (red, Bax+ cells co-expressing the plasma membrane marker Lyn-tdTomato) in a control (left) and an α-catenin deficient embryo (α-catenin MO, right) obtained from the Tg(actb2:Lifeact-GFP) line. Time is indicated in h:min:s. Scale bar: 40 µm.

**Movie 19.** Phagocytic clearance of apoptotic targets (co-expressing the PM marker Lyn-tdTomato, red) in control (left) and α-cateninΔABS overexpressing (right) embryos expressing Lifeact-GFP (cyan). The upper panel shows the phagocytic uptake of apoptotic targets by the embryo epithelium. The lower panel shows phagocytic clearance dynamics within the embryo interior. Embryos were imaged for 3 hours from 30% epiboly (4.7 hpf) to 75% epiboly (8 hpf) and were obtained from the Tg(actb2:Lifeact-GFP) line. Time is indicated in h:min:s. Scale bar: 40 µm.

**Movie 20.** Transient accumulation of Myosin II during the uptake of an apoptotic cell fragment (co-expressing the PM marker Lyn-tdTomato, grey) by epithelial cells. The upper panel shows a top view and the bottom panel shows a transversal view of epithelial cells. Embryos were obtained from the Tg(actb2:myl12.1-EGFP) line and imaged at 50% epiboly (5 hpf). Arrowheads indicate a local enrichment of Myosin II during the uptake of an apoptotic cell. Time is indicated in h:min:s. Scale bars: x: 10 µm, z: 5 µm.

**Movie 21.** Localization of Myosin II at the apical domain and the basal epithelial surface (middle) at the contact sites with apoptotic cells (right) in control embryos. Apoptotic targets (apoptotic Bax+ cells) express a plasma membrane marker (grey, Lyn-tdTomato). Arrowheads indicate the transient accumulation of Myosin II during phagocytic uptake, while asterisks indicate the resulting phagosomes. Embryos were imaged for 1 hour, from 30% epiboly (4.7 hpf) to 50% epiboly stage (5.5 hpf). Embryos were obtained from the Tg(actb2:myl12.1-EGFP) line. Time is indicated in h:min:s. Scale bar: 20 µm.

**Movie 22.** Localization of Myosin II at the apical domain (left) versus its accumulation on the basal epithelial surface (middle) at contact sites with apoptotic cells (right) in E-cadherin morphant (MO) embryos. Apoptotic targets (apoptotic Bax+ cells) express a plasma membrane marker (grey, Lyn-tdTomato). Arrowheads indicate sites of local Myosin II accumulation in epithelial cells, corresponding to contact sites with apoptotic targets. Embryos were imaged for 1h, from 30% epiboly (4.7 hpf) to 50% epiboly stage (5.5 hpf). Embryos were obtained from the Tg(actb2:myl12.1-EGFP) line. Time is indicated in h:min:s. Scale bar: 20 µm.

**Movie 23.** Phagocytic clearance of apoptotic targets (co-expressing the PM marker Lyn-tdTomato, red) in E-cadherin morphant (MO, left) and E-cadherin MO embryos overexpressing the Myosin phosphatase (MP) complex (right), both embryos ubiquitously expressing Lifeact-GFP (cyan). The upper panel shows the phagocytic uptake of apoptotic targets by the embryo epithelium. The lower panel shows the phagocytic clearance dynamics within the embryo interior. Embryos were imaged for 3 hours from 30% epiboly (4.7 hpf) to 75% epiboly (8 hpf) and were obtained from the Tg(actb2:Lifeact-GFP) line. Time is indicated in h:min:s. Scale bar: 40 µm.

**Movie 24.** Localization of GFP-p120-catenin during the uptake of an apoptotic cell fragment (co-expressing the PM marker Lyn-Cerulean, grey) by an epithelial cell. The upper panel shows a top view and the bottom panel shows a transversal view of epithelial cells. Embryos were obtained from the Tg(actb2:Lyn-tdTomato) line injected with GFP-p120-catenin and imaged at 50% epiboly (5 hpf). Arrowheads indicate the local enrichment of GFP-p120-catenin during the uptake of an apoptotic cell fragment. Time is indicated in h:min:s. Scale bars: x: 10 µm, z: 5 µm.

**Movie 25.** Localization of Myosin II at the apical domain (left) versus its accumulation on the basal epithelial surface (middle) at the contact sites with apoptotic cells (right) in p120-catenin MO embryos. Apoptotic targets (apoptotic Bax+ cells) express a plasma membrane marker (grey, Lyn-tdTomato). Arrowheads indicate sites of local Myosin II accumulation corresponding to epithelial interaction sites with apoptotic targets. Embryos were imaged for 1 hour from 30% epiboly (4.7 hpf) to 50% epiboly stage (5.5 hpf). Embryos were obtained from the Tg(actb2:myl12.1-EGFP) line. Time is indicated in h:min:s. Scale bar: 20 µm.

**Movie 26.** Phagocytic clearance in a control embryo (left) and a p120-catenin morpholino injected embryo (right) expressing Lifeact-GFP (cyan). Apoptotic targets (Bax+ cells) express a plasma membrane marker (red, Lyn-tdTomato). The phagocytic uptake of apoptotic targets by the embryo epithelium (upper panel) and the phagocytic clearance dynamics in the embryo interior underneath the epithelium for a control embryo ($z$ = -12 µm) and a p120-catenin MO embryo ($z$ = -6 µm) are shown. Embryos were imaged for 2 hours from 30% epiboly (4.7 hpf) to shield stage (6.5 hpf). Embryos were obtained from the Tg(actb2:Lifeact-GFP) line. Time is indicated in h:min:s:ms. Scale bar: 50 µm.

**Movie 27.** Localization of Myosin II at the apical domain (top) and the basal epithelial surface (middle) at the contact sites with apoptotic cells (Bax+ cells co-expressing the PM marker Lyn-tdTomato, right) in α-catenin MO embryos. Embryos were imaged for 40 min starting from 30% epiboly (4.7 hpf). Embryos were obtained from the Tg(actb2:myl12.1-EGFP) line. Time is indicated in h:min:s. Scale bar: 20 µm.
